# TENT4A non-canonical poly(A) polymerase regulates DNA-damage tolerance via multiple pathways which are mutated in endometrial cancer

**DOI:** 10.1101/2020.12.03.409367

**Authors:** Umakanta Swain, Gilgi Friedlander, Urmila Sehrawat, Avital Sarusi-Portuguez, Ron Rotkopf, Charlotte Ebert, Tamar Paz-Elizur, Rivka Dikstein, Thomas Carell, Nicholas Geacintov, Zvi Livneh

**Author notes:** To whom correspondence should be addressed., Address: Zvi Livneh, Dept. of Biomolecular Sciences, Weizmann Institute of Science, Rehovot, 7610001, Israel.

## Abstract

TENT4A (PAPD7) is a non-canonical poly(A) polymerase, of which little is known. Here we show that TENT4A regulates multiple biological pathways, and focus on its multilayer regulation of translesion DNA synthesis (TLS), in which unrepaired DNA lesions are bypassed by error-prone DNA polymerases. We show that TENT4A regulates mRNA stability and/or translation of DNA polymerase η and RAD18 E3 ligase, which guides the polymerase to replication stalling sites, and monoubiquitinates PCNA, thereby enabling recruitment of error-prone DNA polymerases to damaged DNA sites. Remarkably, in addition to the effect on RAD18 mRNA stability via controlling its poly(A) tail, TENT4A indirectly regulates RAD18 via the tumor suppressor CYLD, and via the long non-coding antisense RNA *PAXIP1-AS2*, which had no known function. Knocking down the expression of *TENT4A* or *CYLD*, or overexpression of *PAXIP1-AS2* led each to reduced amounts of the RAD18 protein and DNA polymerase η, leading to reduced TLS, highlighting *PAXIP1-AS2* as a new TLS regulator. Bioinformatics analysis revealed that TLS error-prone DNA polymerase genes and their *TENT4A*-related regulators are frequently mutated in endometrial cancer genomes, suggesting that TLS is dysregulated in this cancer.

## Introduction

Maintaining genome integrity is critical for the proper functions of cells, and if compromised may lead to severe malfunction and a variety of diseases including cancer, immune malfunction, and neuronal disorders (Akbari, Morevati et al., 2015, Errol, Roger et al., 2006, Hanahan & Weinberg, 2011, Li, Woo et al., 2004, Marteijn, Lans et al., 2014, Modrich, 1994). The majority of DNA lesions inflicted on DNA are removed by accurate DNA repair mechanisms that restore the original DNA sequence. However, the high rate at which DNA lesions are formed (estimated to be thousands/cell/day), and the difficulty in locating them in the genome, makes encounters of DNA replication with DNA lesions inevitable in essentially every cell cycle. Such encounters cause arrest of the replication fork, and if not resolved may lead to fork collapse, double-strand breaks (DSBs), and subsequent chromosomal rearrangements or even cell death. DNA damage tolerance (DDT) mechanisms function in these situations to bypass the lesions, without removing them from DNA, a process that preserved the double-stranded continuity of DNA, giving later a chance to accurate excision repair mechanism to remove the lesion (Branzei & Foiani, 2010, Friedberg, 2005, Livneh, Cohen et al., 2016).

Translesion DNA synthesis (TLS), is a DDT mechanism in which specialized low-fidelity DNA polymerases synthesize across the lesion, a process that is inherently mutagenic (Sale, Lehmann et al., 2012). Despite its mutagenic nature, TLS is surprisingly accurate when replicating across several common DNA lesions such as the sunlight-induced cyclobutane pyrimidine dimers (CPD), and the tobacco-smoke induced (+)-trans-BPDE-N2-dG (BP-G) adduct. To explain this, we have previously proposed that the existence of multiple TLS DNA polymerases enables specialization of certain polymerases to their cognate lesions (meaning more effective and more accurate bypass), and when subjected to the appropriate regulation to ensure the activity of the right polymerase at the right lesion and the right time, this will lead to more accurate TLS, and a lower mutation burden (Cohen, Bar et al., 2015, Hendel, Ziv et al., 2008, Livneh, 2006). We also reported, that p53, and its target gene p21 (acting via its interaction with PCNA), are needed to maintain accurate TLS, partially via their effect on monoubiquitination of PCNA (Avkin, Sevilya et al., 2006). The latter is an important regulatory step, which functions to recruit TLS DNA polymerases to the damaged site in DNA (Hendel, Krijger et al., 2011, Hoege, Pfander et al., 2002, Kannouche, Wing et al., 2004). The molecular deciphering of the regulatory mechanisms of TLS are critical to the understanding of mutation formation, and is the focus of the current study.

*TENT4A* (terminal nucleotidyltransferase 4A; formerly *PAPD7*), is a gene encoding a non-canonical poly(A) RNA polymerase, that we identified in an siRNA-based screen to be required for efficient TLS (Ziv, Zeisel et al., 2014). *TRF4*, its *S. cerevisiae* homolog, was thought to encode a novel DNA polymerase, termed *POLS* (or erroneously, *POLK*), involved in sister chromatid cohesion (Haracska, Johnson et al., 2005, Wang, Castano et al., 2000). However, further studies showed that the *TRF4* and *TRF5* genes encode non-canonical poly(A) RNA polymerases (Haracska et al., 2005). Enzymes of this family are critical in the regulation and quality control of gene expression. There is scarce information about the mammalian *TENT4A*. However, it was reported that its main isoform is 94kDa, considerably larger than the yeast Trf4 and Trf5 proteins (66 and 74 kDa, respectively), and larger than its previously reported size of 62 kDa (Ogami, Cho et al., 2013). Recently TENT4A was shown along with its paralog TENT4B (PAPD5) to synthesize mixed poly(A) tails that contain other inserted nucleotides, primarily guanines, that protect mRNA from deadenylation (Lim, Kim et al., 2018). Still, the functional involvement of TENT4A in biological regulatory pathways is largely unknown (Mueller, Lopez et al., 2019).

Here we report that TENT4A regulates a broad range of genes belonging to a variety of pathways, and focusing on its involvement in TLS we found that *TENT4A* regulates mRNA stability and/or translation of DNA polymerase η (polη ; POLH), and of the RAD18 E3 ligase. Remarkably, in addition to the effect on RAD18-mRNA stability via controlling its poly(A) tail, TENT4A indirectly regulates RAD18 and polη via the tumor suppressor CYLD, and via the long non-coding antisense RNA *PAXIP1-AS2*, which had no known function, highlighting it as a novel TLS regulator. We also partially purified TENT4A and show that it is a poly(A) RNA polymerase, with no detected DNA polymerase activity. Finally, we report that bioinformatics analysis of mutations in the genomes of 33 cancer types from the TCGA database revealed that components of TENT4A-regulated TLS are frequently mutated in endometrial cancer, suggesting involvement of dysregulated TLS in the development of this type of cancer.

## Results

### TENT4A is needed for effective TLS

A screen performed in our lab, identified *TENT4A* (*PAPD7*; *POLS*) as a novel TLS gene (Ziv et al., 2014). To further confirm its effect on TLS, we used the gap-lesion plasmid-based assay, with DNA constructs each containing a single defined lesion (Hendel et al., 2011, Shachar, Ziv et al., 2009, Ziv et al., 2014), and two cell lines: The human osteosarcoma U2OS cell line and the MCF-7 breast cancer cell line. As can be seen in Fig 1A, B and C, Appendix Fig S1, and Appendix Tables S1 and S4, knocking down the expression of *TENT4A* in these cells, but not of *TENT4B*, led to a decrease in TLS across four types of DNA lesions: a BP-G adduct, a major lesion caused by tobacco smoke; a thymine-thymine cyclobutane pyrimidine dimer (TT-CPD) and a thymine-thymine 6-4 photoproduct (TT-6-4PP), the most common, and the second most common type of UV light-induced DNA damage, respectively; and a cisplatin guanine-guanine (cisPt-GG), a major intrastrand crosslink formed by the chemotherapeutic drug cisplatin. DNA sequence analysis showed that the error frequency of the residual TLS under *TENT4A* knockdown did not substantially change (Appendix Tables S2, S3, S5 and S6). The broad DNA damage spectrum of the effect suggests that *TENT4A* regulation of TLS is not directed to a specific TLS DNA polymerase, but instead functions in a rather more general TLS regulatory step that exerts a global effect.

**Figure 1.**
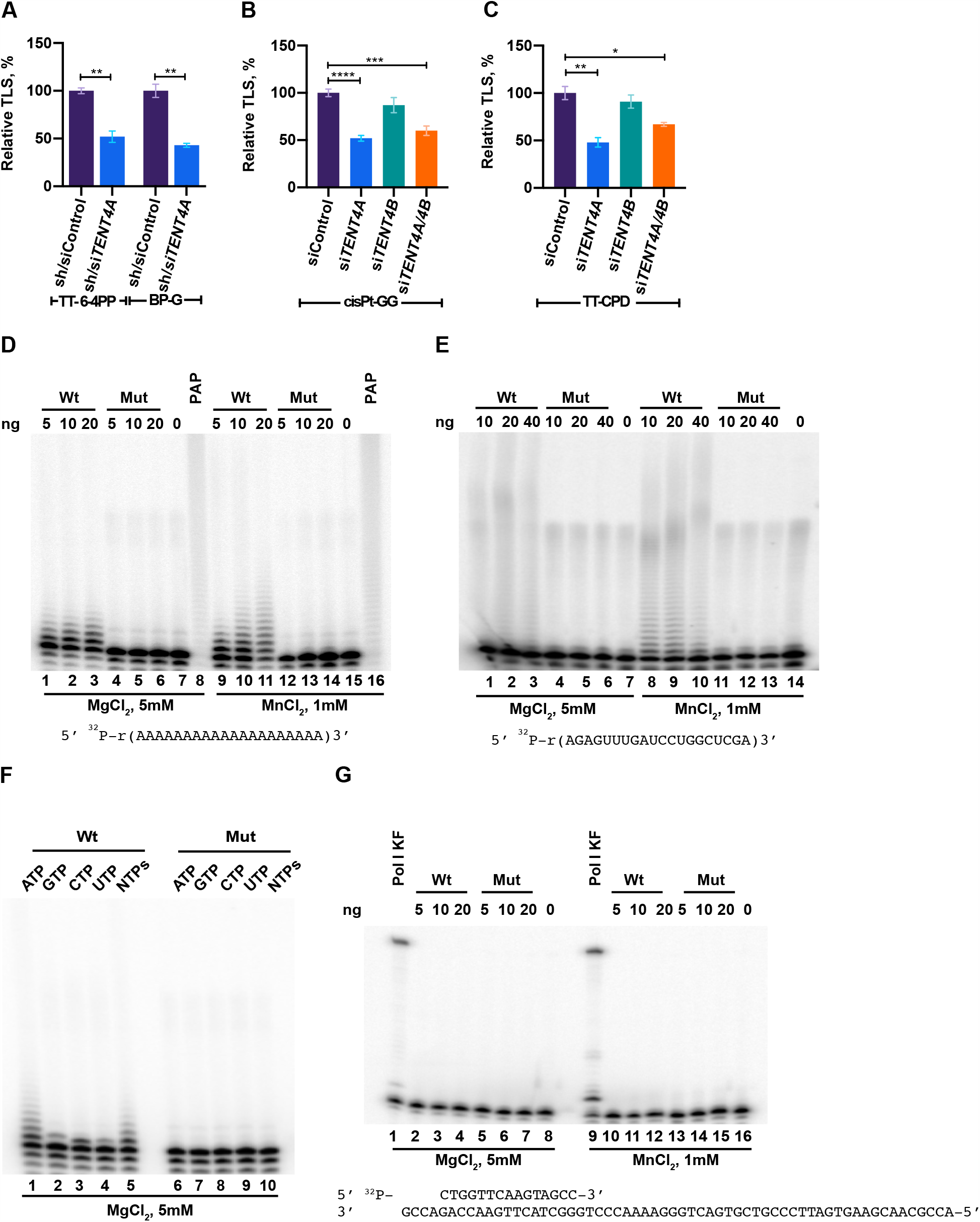
Involvement of *TENT4A* in TLS, and its poly (A) RNA polymerase enzyme activity. **(A)** TLS across a TT-6-4PP and BP-G in U2OS cells. *TENT4A* expression was knocked-down using lentivirus-mediated shRNA combined with siRNA against *TENT4A*. The results are presented as the mean ± SEM of three independent experiments (see Appendix Table S1 for details). Statistical analysis was performed using the two-tailed Student’s *t*-test (\*\**P* < 0.01). **(B)** and **(C)** TLS across a cisPt-GG and TT-CPD, respectively, in MCF-7 cells in which the expression of *TENT4A* and/or *TENT4B* genes was knocked-down using specific siRNAs. The results are presented as the mean ± SEM of six independent experiments for cisPt-GG and three independent experiments for TT-CPD (see Appendix Table S4 for details). Statistical analysis was performed using the two tailed Student’s *t*-test (\*\*\*\**P* < 0.0001, \*\*\**P* < 0.001, \*\**P* < 0.01, \**P* < 0.05). **(D)** Poly(A) RNA polymerase activity measured by the extension of an oligo (A)_20_ substrate. Poly(A) polymerase assays were performed with the indicated amount of partially purified recombinant human 8xHis-MBP tagged TENT4A (lanes 1-3 and 9-11) or the TENT4A mutant D277A, D279A (lanes 4-6 and 12-14), using 5’ ^32^P-labeled oligo(A)_20_ and 1 mM ATP in presence of 5 mM MgCl_2_ (lanes 1-6) or 1 mM MnCl_2_ (lanes 9-14). For positive and negative controls, parallel reactions were also carried out with 0.5 units of *E. coli* poly(A) polymerase (lanes 8 and 16) or no added protein (lanes 7 and 15), respectively. The 5’ ^32^P-labeled RNA oligo sequence is shown below the image. Reaction products were resolved on a 15% polyacrylamide gel containing 8 M urea and analyzed by PhosphorImaging. **(E)** Poly(A) RNA polymerase assays with an oligo RNA. The assays were performed as described in panel **(D)**, except that the oligo RNA shown underneath the gel image was used, as well as the four NTPs (1 mM each). **(F)** Ribonucleotide specificity of TENT4A. The ^32^P-labeled RNA oligo(A)_20_ substrate was incubated with the TENT4A protein (lanes 1-5) or its mutant (lanes 6-10) in the presence of each of the four NTPs (1 mM each). **(G)** Assays of TENT4A DNA polymerase activity. Assays were performed with the indicated amount of partially purified recombinant human 8xHis-MBP tagged TENT4A (lanes 2-4 and 10-12) and mutant (lanes 5-7 and 13-15), using 0.5 pmol of ^32^P-labeled 15/60-nt primer/template in the presence of the four dNTPs (100 µM each), 5 mM MgCl_2_ (lanes 2-7) or 1 mM MnCl_2_ (lanes 10-15). For positive and negative controls, parallel reactions were carried out with the Klenow fragment of Pol I (lanes 1 and 9), or no added protein (lanes 8 and 16), respectively. The sequence of the primer/template DNA substrate is shown below the image.

### Purified TENT4A is a non-canonical poly(A) RNA polymerase, which incorporates also other ribonucleotides

We partially purified overexpressed full-length human TENT4A as an 8xHis-MBP-tagged protein, (Appendix Fig S2), and assayed its potential activities as a poly(A) RNA polymerase and DNA polymerase. In parallel, we purified a TENT4A variant that carries the D277A and D279A mutations (Appendix Fig S2), which were expected to inactivate the coordination of the divalent metal ion as a co-substrate. As can be seen in Fig 1D, TENT4A used ATP in the presence of Mg^++^ to extend the oligo (A)_20_ substrate. In contrast, the TENT4A mutant was inactive. A similar activity was observed with the oligo 5’ r(AGAGUUUGAUCCUGGCUCGA)-3’ (Fig 1E). Unlike *E. coli* poly(A) RNA polymerase, which was used as a positive control, and performed extensive extension of the substrate, TENT4A added a limited number of AMP residues, which is typical of a non-canonical poly(A) RNA polymerase activity (Fig 1D and E). The polyadenylation activity of TENT4A was stimulated when Mn^++^ was used instead of Mg^++^, whereas the mutant TENT4A remained unaffected (Fig 1D and E). TENT4A preferentially polymerizes AMP residues, although it can use other rNTPs to some extent (Fig 1F), consistent with a previous report (Lim et al., 2018). Of note, the shorter form of TENT4A, previously believed to be the full-length enzyme, had negligible poly(A) RNA polymerase activity (not shown). We also examined the DNA polymerase activity of the recombinant tagged TENT4A, and found no activity (Fig 1G). We conclude that TENT4A is a poly(A) RNA polymerase, and as far as we can tell, it has no detectable DNA polymerase activity.

### Effect of *TENT4A* on mRNA of genes involved in TLS

Poly(A) RNA polymerases are typically involved in regulating mRNA via 3’-polyadenylation, and therefore to start addressing the mechanism by which *TENT4A* regulates TLS, we examined whether the mRNA of TLS-related genes bind TENT4A. To that end, we used RNA immunoprecipitation-qPCR (RIP-qPCR). We overexpressed FLAG-TENT4A, then immunoprecipitated the protein using anti-FLAG antibody, and extracted the RNA from protein-RNA complexes. The amount of specific mRNA was determined by preparing cDNA, and performing qPCR using gene specific primers. As can be seen in Fig 2A, while the mRNAs of *POLH, POLK* and *POLI* showed no significant binding, the mRNAs of *REV3L* encoding the catalytic subunit of DNA polymerase ζ, and of *REV1*, encoding a TLS scaffold protein/polymerase, as well as *RAD18* and *CYLD*, showed significant preferential binding. RAD18 is the E3 ligase that monoubiquitinates PCNA (Watanabe, Tateishi et al., 2004), a central signalling event in TLS, and CYLD is a deubiquitinase involved in cancer and required for efficient TLS, as we have previously reported (Ziv et al., 2014).

**Figure 2.**
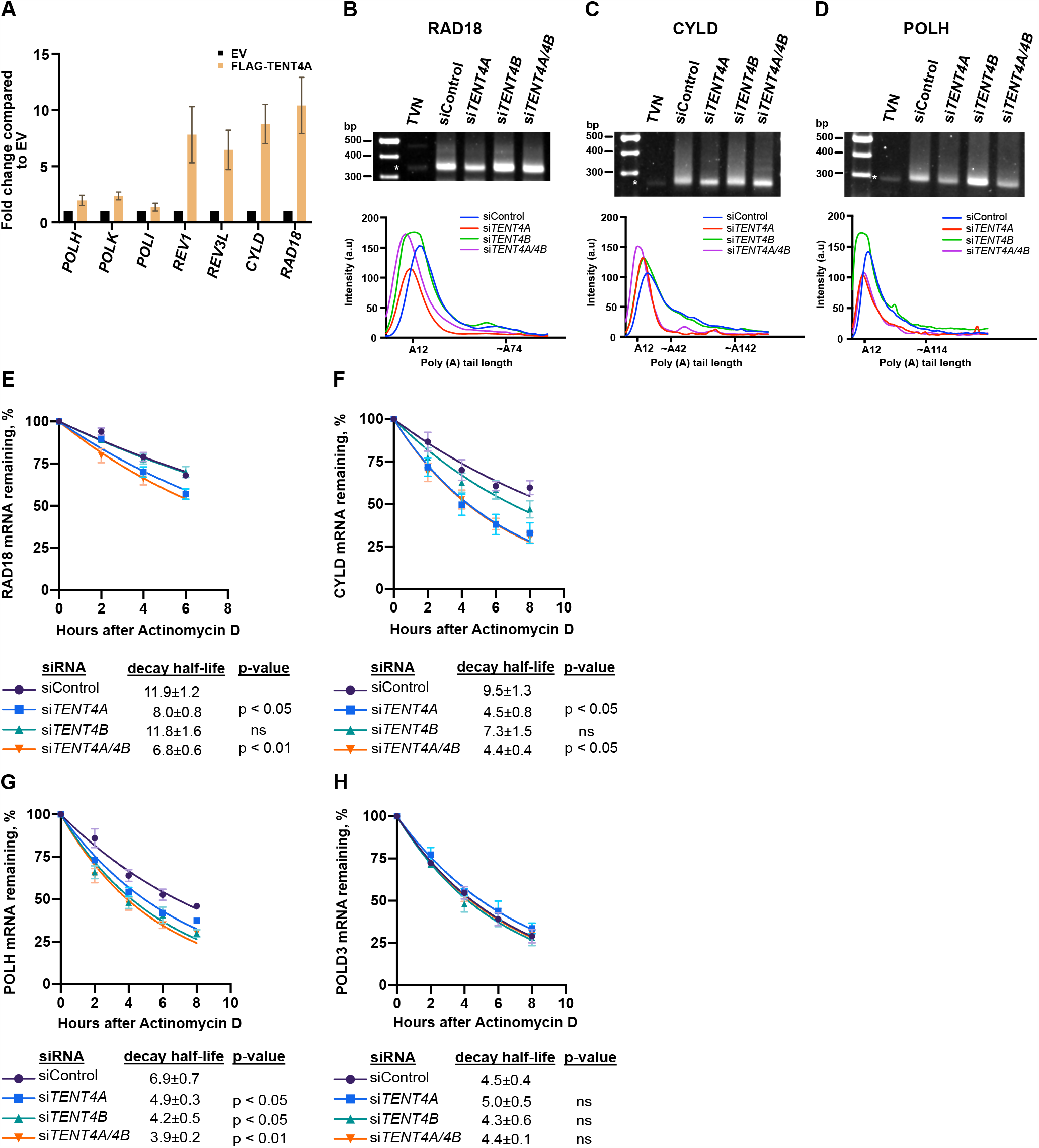
Effect of TENT4A on binding and stability of TLS-related mRNAs. **(A)** FLAG-tagged TENT4A expressed in HEK293T cells was immunoprecipitated with an anti-FLAG antibody, and RNA bound to it was analyzed by qPCR. The results are shown as fold-change relative to empty vector. The error bars indicate SEM (two independent experiments). **(B), (C)**, and **(D)** Representative gel images of *RAD18, CYLD*, and *POLH* poly(A) tail are shown, each with the poly(A) tail densitometric trace underneath, using Image J. The position of TVN is indicated by an oligonucleotide that predominantly recognizes A_12_, serving as an internal control. **(E), (F), (G), (H)**, *TENT4A* and/or *TENT4B* expression was knocked-down in MCF-7 cells for 48 h, after which the cells were treated with 5 μg/ml Actinomycin D for up to 8 h. The expression levels of *RAD18, CYLD, POLH*, and *POLD3* genes respectively, were each determined by qPCR relative to the expression level at time 0. Half-life was calculated by using non-linear fit one phase decay curve equation and significance of the differences was calculated using Student’s *t*-test (one-tailed); Data are presented as mean ± SEM from three independent experiments. ns: *P*>0.05.

To examine whether TENT4A affects the length of the poly(A) tail of these genes, we used ePAT (extension poly(A) test; (Janicke, Vancuylenberg et al., 2012)). We examined the TLS regulators *RAD18* and *CYLD*, which bound TENT4A, and also *POLH*, which did not bind TENT4A. As can be seen in Fig 2, knocking down the expression of *TENT4A* caused a shift to shorter amplified tail fragments of *RAD18*, and similar results were obtained for the tail fragments of *CYLD* (Fig 2B and C). Interestingly, knocking down the expression of *TENT4A* caused a shift to shorter amplified tail fragments of *POLH* too, despite its lack of binding to TENT4A, suggesting indirect regulation. Knocking down both *TENT4A* and *TENT4B* caused a similar shift (Fig 2B, C and D).

Because the length of the poly(A) tail may affect mRNA stability, we examined the effect of *TENT4A* on the stability of the mRNA of these TLS-related genes by measuring their half-life in the presence of the transcription inhibitor Actinomycin D. As can be seen in Fig 2E, knocking down the expression of *TENT4A*, but not *TENT4B*, caused a decrease of about 33% in the half-life of *RAD18* mRNA, whereas knocking down the expression of both caused a slightly stronger decrease of 43%. The half-life of *CYLD* mRNA was decreased by 53% when the expression of *TENT4A* was knocked-down, with a smaller effect of *TENT4B* knockdown (Fig 2F). Interestingly, for *POLH* mRNA, knocking down the expression of *TENT4A* had a decrease in half-life (29%), slightly lower than the effect of *TENT4B* knockdown (39% decrease) (Fig 2G). The stability of a control *POLD3* mRNA was essentially unaffected by knocking down either *TENT4A, TENT4B* or both (Fig 2H), as was the stable *RNA18S* rRNA (Appendix Fig S3). These results suggest that *TENT4A* directly regulates the stability of *RAD18* and *CYLD* mRNA, and indirectly the stability of *POLH* mRNA via the length of their poly(A) tails.

### *TENT4A* affects the amounts of RAD18, CYLD and POLH proteins via different mechanisms

Since effects on mRNA are often, but not always, manifested in the levels of the encoded proteins, we next analysed whether *TENT4A* affects also the protein products of the *RAD18, POLH* and *CYLD* genes. As can be seen in Fig 3A, upon knocking down the expression of *TENT4A*, there was a decrease in the amount of RAD18, observed in both unirradiated and UV-irradiated cells. Knocking down *TENT4A* expression led also to a decrease in the amounts of the CYLD protein (Fig 3B) and polη (*POLH* gene product (Fig 3B).

**Figure 3.**
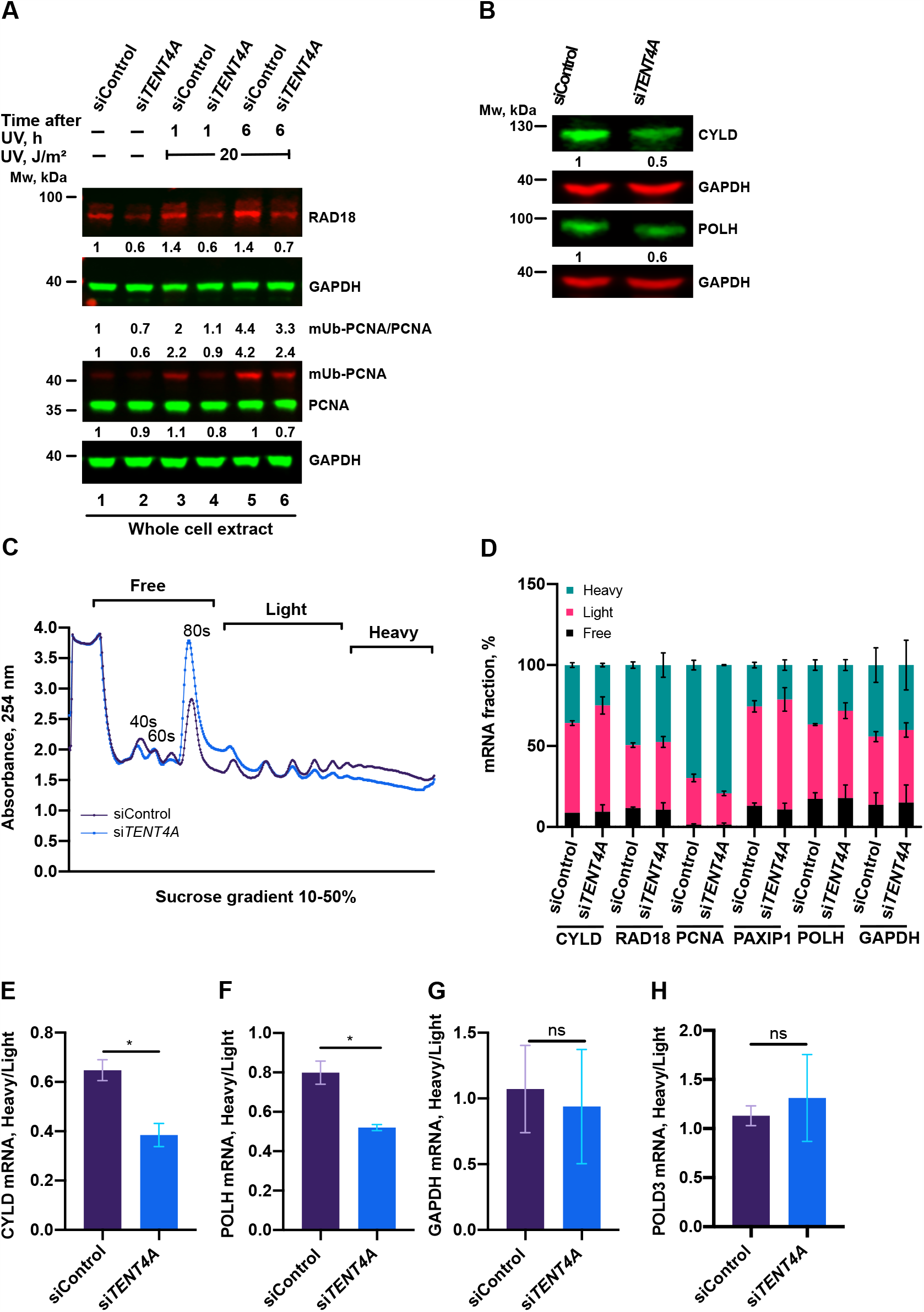
Effect of *TENT4A* on levels and translation of TLS proteins. **(A)** MCF-7 cells were transfected with *TENT4A*-targeted siRNA (si*TENT4A*) or non-targeting control siRNA (siControl). At 65 h post-transfection, the cells were UV-irradiated at 20 J/m^2^ and harvested 1 or 6 h post-irradiation. Whole cell extracts were fractionated by SDS-PAGE followed by western blot with indicated antibodies. Protein levels were normalized to GAPDH, and presented each relative to its level in unirradiated siControl-treated cells. **(B)** Effect of cell treatment (72 h) with si*TENT4A* or siControl on the levels of the CYLD and POLH proteins in MCF-7 cells. Whole cell extracts were fractionated by SDS-PAGE and analyzed by western blot with indicated antibodies. Protein levels were normalized to GAPDH, and presented each relative to its level in siControl-treated cells. **(C)** Polysome profile analysis in MCF-7 cells in which *TENT4A* expression was knocked-down. Absorbance trace of polysome fractionation on a sucrose gradient. **(D)** qPCR analysis of mRNA levels of TLS-related genes in each fraction were expressed as a percentage of total levels summed over all fractions. The results are presented as the mean ± SEM of two independent experiments. **(E), (F)** and **(G)** Fraction of polysome-associated mRNA of the genes *CYLD, POLH* and *GAPDH*, respectively. The values were taken from **(D)**. (**H**) Fraction of polysome-associated mRNA of the *POLD3* control gene. The results are presented as the mean ± SEM of two independent experiments. Statistical significance was determined using Student’s *t*-test (one-tailed); \**P* < 0.05, ns: *P* > 0.05.

To explore whether this reduction in the amounts of the RAD18, CYLD and POLH proteins was a result of reduced translation efficiency, we performed polysome profiling under condition in which *TENT4A* expression was knocked-down. As can be seen in Fig 3C, there was an increase in the 80S monosome and a decrease in the polysomal fractions when *TENT4A* expression was knocked-down, indicating an inhibition of translation efficiency. Analysis of individual genes revealed a shift from the heavy to the light fractions of *CYLD* and *POLH* mRNA (Fig 3E and F), but not of *RAD18* mRNA (Fig 3D). Translation of the control gene *POLD3* (Fig 3H) was unaffected by knocking down the expression of *TENT4A*. Thus, the reduction in CYLD and POLH protein amount following *TENT4A* knockdown is due to both their reduced mRNA stability and its reduced translation efficiency, but the decreased amount of RAD18 protein following *TENT4A* knockdown is due to reduced mRNA stability, as the translation efficiency was unaffected.

### Regulation of PCNA monoubiquitination by TENT4A

RAD18 is the key E3 ligase that monoubiquitinates PCNA, and we therefore examined whether the *TENT4A* effect of reducing the amounts of RAD18 is also manifested in the level of mUb-PCNA. As can be seen in Fig 4, upon *TENT4A* knockdown the amount of mUb-PCNA was mildly decreased in UV-irradiated cells (Fig 4, lane 10), and strongly decreased in unirradiated cells (Fig 4, lane 2). The amount of mUb-PCNA was calculated as the mUb-PCNA/PCNA ratio, and as the amount (independent of PCNA) normalized to the loading control GAPDH, generally showing a similar behaviour upon *TENT4A* knockdown.

**Figure 4.**
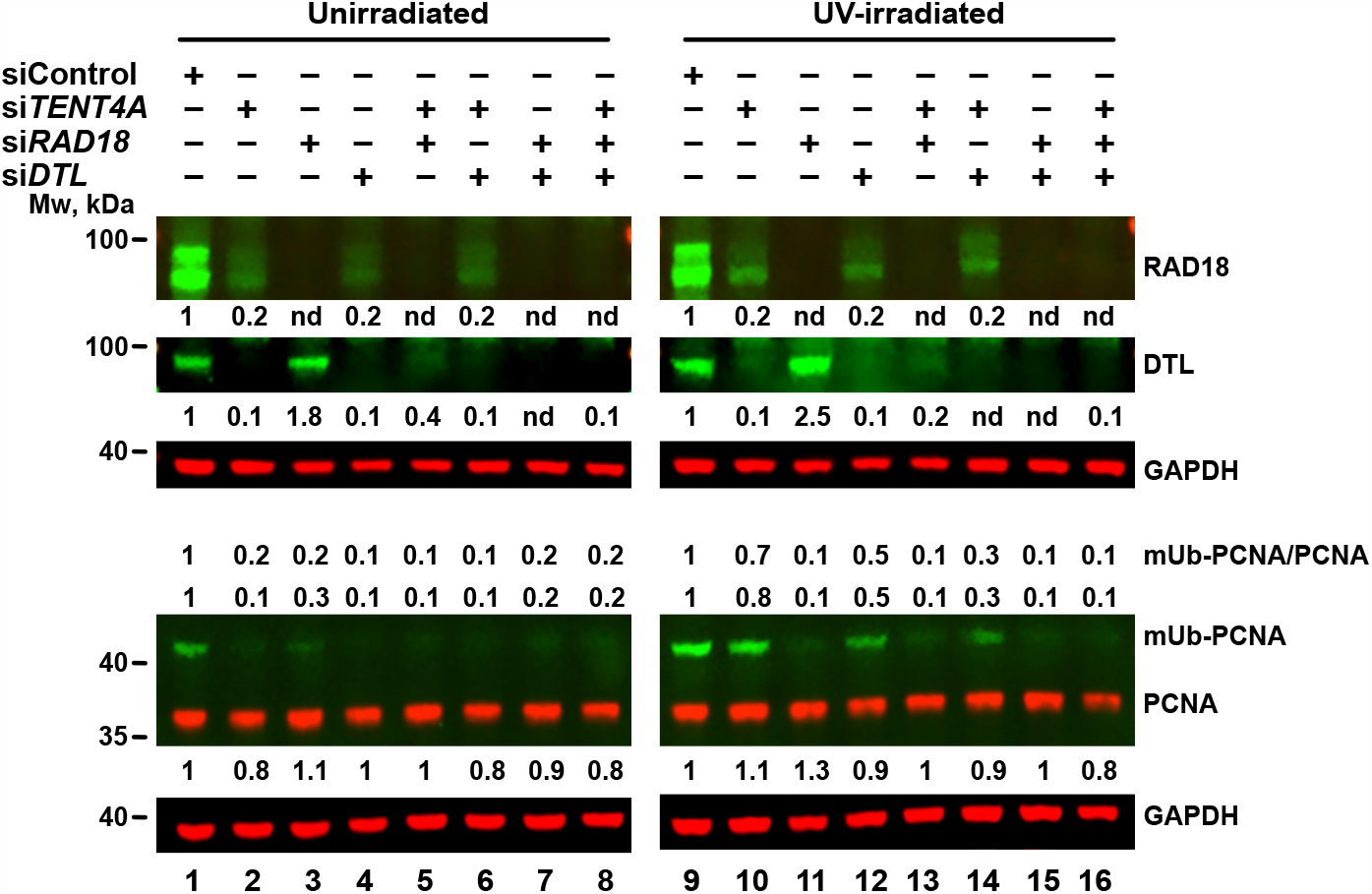
Interrelationship among TENT4A, RAD18 and DTL in determining amounts of RAD18 and DTL and the extent of PCNA monoubiquitination. MCF-7 cells were transfected with siRNA against *TENT4A, RAD18*, and *DTL* or their double or triple combinations. At 65 h post-transfection, the cells were UV-irradiated at 20 J/m^2^ and harvested 6 h post-irradiation. Whole cell extracts were fractionated by SDS-PAGE followed by western blot with indicated antibodies. Protein levels were normalized to GAPDH and presented each relative to its level in unirradiated or UV-irradiated siControl-treated cells.

PCNA monoubiquitination in untreated cells was reported to be carried out by the DTL (CTD2) E3 *ligase* (Terai, Abbas et al., 2010), and we therefore examined the effect of *TENT4A* knockdown on DTL. As can be seen in Fig 4 lanes 2 and 10, knocking down the expression of TENT4A caused a strong reduction in the amount of the DTL protein, in both unirradiated and UV-irradiated cells. In parallel, also the amount of the RAD18 protein was decreased, consistent with the results shown above. We then examined whether RAD18 and DTL affect each other. Interestingly, when the expression of *DTL* was knocked-down, the amount of the RAD18 protein was also reduced in both unirradiated (Fig 4, lane 4) and UV-irradiated cells (Fig 4, lane 12). On the other hand, when the expression of RAD18 alone was knocked-down, the amount of DTL was *increased*, in both unirradiated (Fig 4, lane 3) and UV-irradiated cells (Fig 4, lane 11). When both *TENT4A* and *RAD18* were knocked-down, DTL was still decreased, but to an extent lesser than with *TENT4A* knockdown alone, indicating an additive effect (Fig 4, lanes 5 and 13). On the other hand, the *TENT4A* and *DTL* double knockdown had a similar effect to the single gene knockdown, indicating the two are epistatic (Fig 4, lanes 6 and 14).

How do these effects on RAD18 and DTL translate into levels of mUb-PCNA? In UV-irradiated cells *RAD18* knockdown caused a strong reduction in mUb-PCNA, despite the increase in DTL, consistent with RAD18 being the major E3 ligase that monoubiquitinates PCNA in UV-irradiated cells, with a similar effect observed with the *TENT4A* and *RAD18* double knockdown (Fig 4, lanes 11 and 13). Interestingly, knocking down DTL caused a mild decrease in mUb-PCNA in UV-irradiated cells (Fig 4, lane 12), likely because under these conditions RAD18 was also decreased. In unirradiated cells, knocking down the expression of DTL caused a strong decrease in mUb-PCNA, consistent with previous work (Terai et al., 2010), but also knocking down RAD18 caused a decrease in mUb-PCNA (Fig 4, lane 3), despite a slight increase in the amount of DTL. Similar effects were observed when both *TENT4A* and *DTL* were knocked-down (Fig 4, lane 6). When both *TENT4A* and *RAD18* were knocked-down, mUb-PCNA was strongly reduced (Fig 4, lane 5), as under these conditions the two E3 ligases were reduced.

### TENT4A-regulated *CYLD* regulates RAD18, DTL and POLH

*CYLD*, previously identified by us in a screen as a TLS regulator, acts downstream to *TENT4A*, and is regulated by *TENT4A* at the mRNA stability and translation levels as shown above. To examine its own effect on TLS, we knocked-down *CYLD* expression, and examined levels of RAD18, mUb-PCNA and POLH. As can be seen in Fig 5A, knocking down *CYLD* expression caused a strong 3-5 fold decrease in the amount of RAD18, clearly visible in both unirradiated and UV-irradiated cells. Similarly, POLH protein decreased about 3-fold (Fig 5A). It thus appears that in addition to its direct effect on RAD18, *TENT4A* has an indirect effect on both RAD18 and POLH proteins, mediated via its target gene *CYLD*.

**Figure 5.**
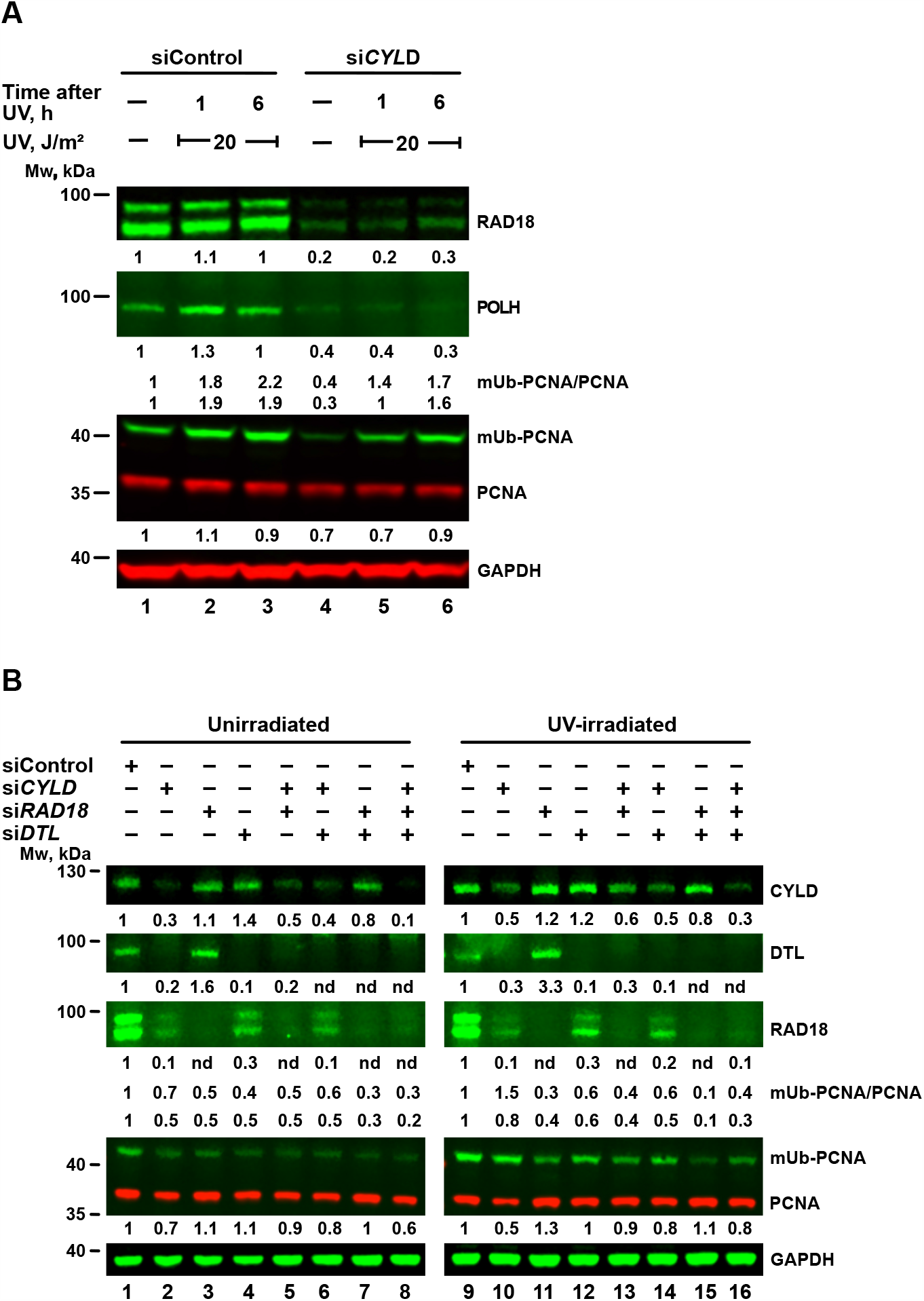
Effect of CYLD on RAD18, DTL, POLH and PCNA monoubiquitination. (**A**) MCF-7 cells were transfected with *CYLD*-targeted siRNA (si*CYLD*) or siControl. At 65 h post-transfection, the cells were irradiated at 20 J/m^2^ UV and harvested 1 or 6 h post-irradiation. (**B**) MCF-7 cells were transfected with siRNA against *CYLD, RAD18*, and *DTL* or their double or triple combinations. At 65 h post-transfection, the cells were UV-irradiated at 20 J/m^2^ and harvested 6 h post-irradiation. Whole cell extracts were fractionated by SDS-PAGE and analyzed by western blot with indicated antibodies. Protein levels were normalized to GAPDH and presented each relative to its level in unirradiated or UV-irradiated siControl-treated cells.

An analysis of mUb-PCNA in cells in which *CYLD* was knocked-down revealed that despite the reduction in RAD18 protein, UV-induced mUb-PCNA levels were little affected (Fig 5A). Assuming that DTL might be backing up RAD18 under these conditions, we knocked-down the expression of *CYLD, RAD18* or *DTL* individually, or in combinations of two or three genes. As can be seen in Fig 5B, knocking down *CYLD* caused a strong decrease of both RAD18 and DTL, in both unirradiated (Fig 5B, lane 2) and UV irradiated cells (Fig 5B, lane 10). Of note, a decrease in RAD18 protein was not always paralleled by a decrease in mUb-PCNA (Fig. 5B).

### *TENT4A* regulates multiple genes and pathways, including long non-coding RNAs and antisense transcripts

Because *TENT4A*, being a poly(A) RNA polymerase, is expected to exert its effect via RNA, we conducted an RNA-seq analysis, comparing genes in control cells to the same cells in which *TENT4A* was knocked-down. The experiments were conducted in triplicates in two different human cell lines, in which the expression of *TENT4A* was knocked-down with shRNA plus siRNA (U2OS), or siRNA (XP12RO). Overall the expression of a large number of genes was altered (Fig 6A and Appendix Data 1), with an intersection of 313 genes downregulated, and 149 genes upregulated in the two cell lines (Fig 6A). A heatmap showed that the overall expression pattern of the shared genes was similar in the two cell lines (Fig 6B).

**Figure 6.**
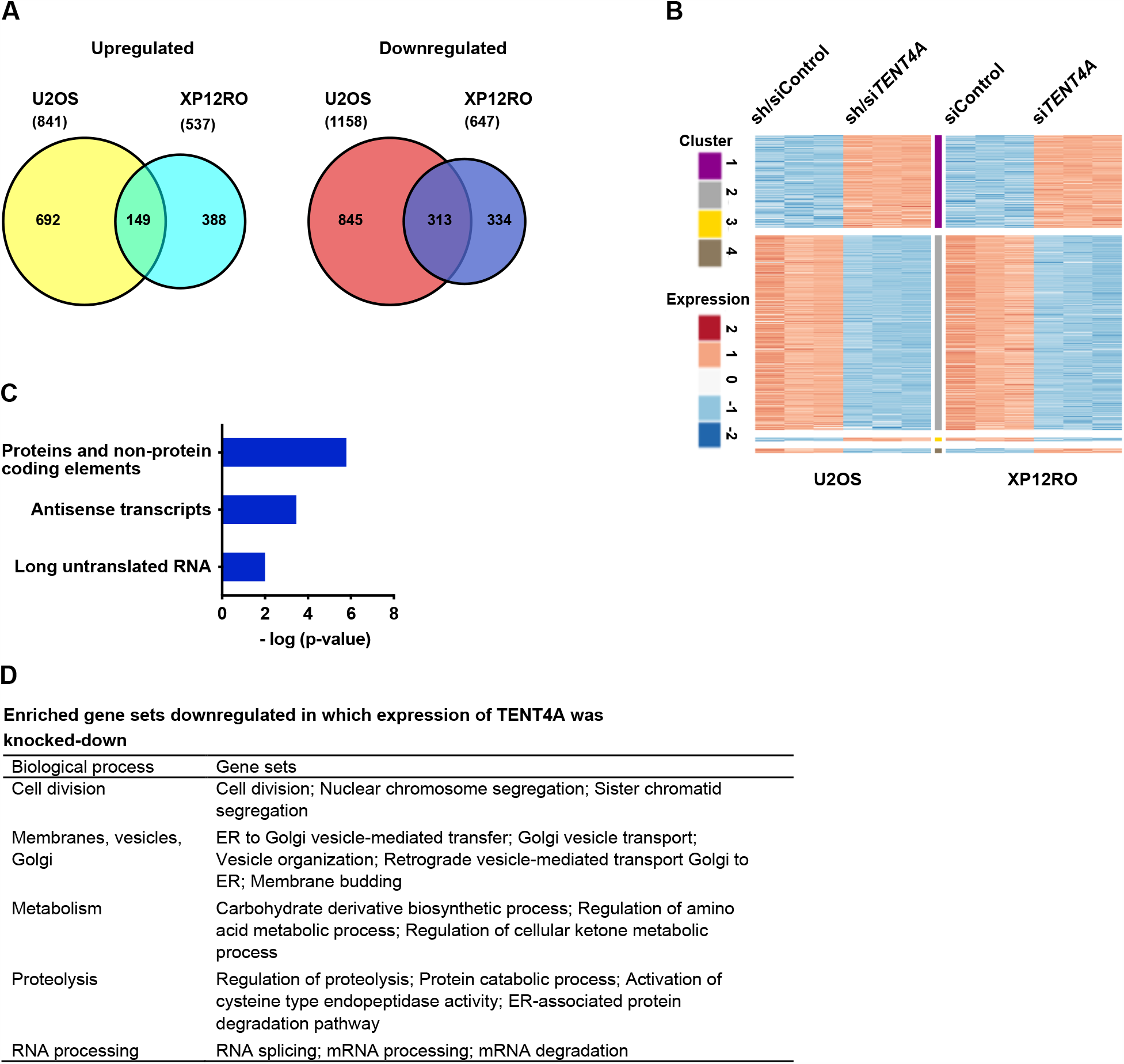
RNA-seq analysis of cells in which the expression of *TENT4A* was knocked-down. **(A)** Venn diagram illustrating the number of significantly differentially expressed genes (FDR ≤ 0.05, absolute fold change ≥ 2 and a count of at least 30 at least in one of the samples) in *TENT4A* knocked-down U2OS and XP12RO cells. The numbers of upregulated, downregulated and overlapping genes are shown (see detailed RNA-seq results in Appendix Data 1). **(B)** Heatmap representation of the shared significantly differentially expressed genes in U2OS and XP12RO experiments (a total of 475 genes). The response of most genes is the same in both cell lines (149 are upregulated and 313 are downregulated). For each experiment, the log_2_ normalized counts were standardized to have for each gene zero mean and unit standard deviation. K-mean clustering of the two data-sets using Euclidean distance measure is presented. The expression profile is accompanied by a colored bar indicating the standardized log_2_ normalized counts. **(C)** Pathway Studio ontology of differentially expressed upregulated genes that are common to U2OS and XP12RO analyzed by Pathway Studio. Ontology details are in Appendix Data 2. **(D)** Gene sets enriched in downregulated genes in *TENT4A* knocked-down U2OS and XP12RO cells. The mRNA degradation pathway was from the Pathway Studio analysis. See details data in Appendix Data 2.

Gene set enrichment analysis (GSEA; Broad Institute) revealed that the significantly downregulated genes in *TENT4A* cells, corresponded to 44 gene sets that were enriched in both cell lines with an FDR of <0.05 (Appendix Data 2). No gene sets with upregulated genes were common for both cell lines. Of note, Pathway Studio analysis revealed among genes whose expression was upregulated, also antisense transcripts and long untranslated RNA (Fig 6C). Overall, the biological pathways with genes that were down-regulated can be categorized to 5 main processes, namely (1) Cell division; (2) Membranes, vesicles and Golgi; (3) Metabolism; (4) Proteolysis; (5) RNA processing; (Fig 6D). Thus, *TENT4A* appears to regulate many genes involved in diverse biological processes, including mRNA degradation, as expected from a poly(A) RNA polymerase, and consistent with its effect on the half-life of several TLS-related mRNAs (Fig 2). Of note, Gene Ontology defined DNA repair pathways were downregulated upon treatment with siRNA against *TENT4A* (Appendix Data 2).

### A *TENT4A*-regulated long non-coding antisense RNA to the *PAXIP1* (*PTIP*) gene regulates RAD18, POLH and mUb-PCNA

As mentioned above, the RNA-seq analysis revealed that knocking down the expression of *TENT4A* causes upregulation of long non-coding RNA and antisense transcripts. One of these is *PAXIP1-AS2*, which is a long non-coding RNA, antisense to the *PAXIP1* gene. Because PAXIP1 was reported to promote PCNA monoubiquitination (Gohler, Munoz et al., 2008), we studied the involvement in TLS of *PAXIP1-AS2*, for which no biological function was yet assigned.

Knocking down the expression of *TENT4A* caused a significant increase in the amount of the *PAXIP1-AS2* transcript (Fig 7A), as expected from the RNA-seq analysis. This effect was not observed when *TENT4B* was knocked-down, suggesting that it is specific to *TENT4A* (Fig 7A). We next overexpressed *PAXIP1-AS2* and examined its effect on TLS. As can be seen in Fig 7B and Appendix Tables S7 and S8, TLS across cisPt-GG was strongly suppressed by overexpressing *PAXIP1-AS2*. This effect was not mediated via the amount or stability of *PAXIP1* mRNA, which remained essentially unchanged upon overexpression of the antisense RNA (Fig 7C and D), however, the amount of PAXIP1 protein was diminished (Fig 7E). Directly knocking down the expression of *PAXIP1* also inhibited TLS across a TT-CPD lesion (Fig 7F and Appendix Tables S9 and S10), consistent with the effect of the antisense RNA.

**Figure 7.**
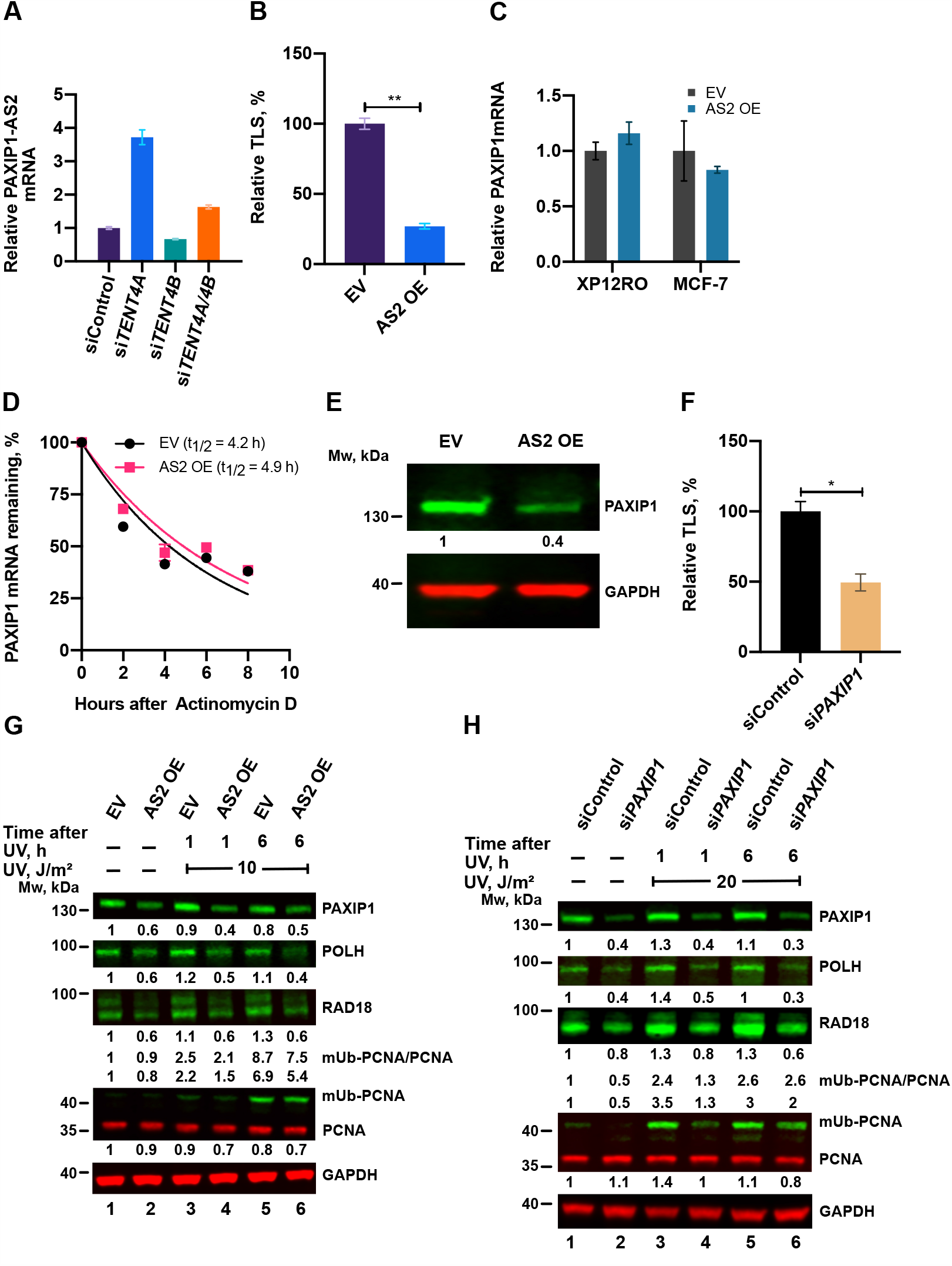
Involvement of *PAXIP1-AS2* and *PAXIP1* in TLS. **(A)** Expression of *TENT4A* and/or *TENT4B* was knocked-down for 48 h, after which the level of *PAXIP1-AS2* transcript was measured by qPCR, normalized to *GAPDH*, and compared to cells treated with siControl. The results are presented as the mean ± SEM of three independent experiments. **(B)** TLS across a cisPt-GG in MCF-7 cells in which *PAXIP1-AS2* antisense RNA was overexpressed (See Appendix Tables S7 and S8). The results are presented as the mean ± SEM of two independent experiments. Statistical analysis was performed using the two-tailed Student’s *t*-test (\*\**P* < 0.01). **(C)** *PAXIP1-AS2* antisense RNA was overexpressed in XP12RO or MCF-7 cells for 48 h, after which the amount of *PAXIP1* mRNA was measured by qPCR, normalized to *GAPDH* and presented relative to cells transfected with an empty vector (EV). The results are presented as the mean ± SEM of three independent experiments. **(D)** *PAXIP1-AS2* antisense transcript was overexpressed for 48 h in MCF-7 cells, after which *PAXIP1* mRNA levels were determined as described in Fig 2. *GAPDH* was used as normalized control. The results are presented as the mean ± SEM of two independent experiments. **(E)** Effect of overexpression of *PAXIP1-AS2* antisense RNA on PAXIP1 protein. Cells were harvested after 48 h and whole cell extracts were analyzed by SDS–PAGE followed by western blot with indicated antibodies. **(F)** TLS across a TT-CPD lesion in MCF-7 cells in which the expression of *PAXIP1* was knocked-down for 68 h. TT-CPD TLS assay was performed for 4 h. See Appendix Tables S9 and S10 for details. The results are presented as the mean ± SEM of two independent experiments. Statistical analysis was performed using the two-tailed Student’s *t*-test (\**P* < 0.05). **(G)** Effect of overexpression of *PAXIP-AS2* on TLS-related proteins in UV-irradiated cells. *PAXIP1-AS2* was overexpressed in XP12RO cells for 48 h after which cells were UV-irradiated at 10 J/m^2^ UV and harvested 1 or 6 h post-irradiation. Whole cell extracts were analyzed by SDS– PAGE followed by western blot with indicated antibodies. Protein levels are presented relative to those in extracts of unirradiated cells transfected with empty vector. **(H)** Effect of *PAXIP1*-knockdown on TLS proteins. MCF-7 cells were knocked-down by siRNA against *PAXIP1* for 72 h, after which cells were UV-irradiated at 20 J/m^2^ UV and harvested 1 or 6 h post-irradiation. Whole cell extracts were analyzed by SDS–PAGE and detected with indicated antibodies. Amounts are presented relative to those in extracts of unirradiated cells treated with siControl.

We next examined the effects of *PAXIP1* and *PAXIP1-AS2* on POLH, RAD18 and mUb-PCNA. As can be seen in Fig 7G and H, *PAXIP1-AS2* overexpression, and similarly knockdown of *PAXIP1*, each caused a decrease in the amount of POLH, whereas the effect on RAD18 and mUb-PCNA was marginal.

As described above, TENT4A did not bind *POLH* mRNA, yet it affected its stability, raising the possibility that the effect was mediated via *PAXIP1*. Knocking down the expression of *PAXIP1* reduced the half-life of *POLH* mRNA to an extent similar to knocking-down the expression of *TENT4A* (Fig 8A). Knocking down both *TENT4A* and *PAXIP1* caused a decrease of *POLH* mRNA stability similar to that of each gene alone, suggesting that the two are epistatic, and that possibly the effect of *TENT4A* on *POLH* mRNA is mediated largely via *PAXIP1*. Western blot analysis revealed that the decrease in polη amount was more pronounced under *PAXIP1* than *TENT4A* knockdown (Fig 8C). Thus, while the effects of the two seem to be epistatic at the mRNA level, additional regulation affects the polη protein level by mechanisms which are yet to be explored, e.g., via a positive regulation by *PAXIP1*, or a branch of negative regulation by *TENT4A*. A similar epistatic analysis was performed with *TENT4A* and *PAXIP1* for *RAD18* mRNA, which binds TENT4A. Knocking down the expression of *TENT4A* had a bigger effect than *PAXIP1* of decreasing the half-life of *RAD18* mRNA, while knocking down the two had an intermediate effect (Fig 8B). A similar effect of observed also at the protein level, in both unirradiated, and UV-irradiated cells (Fig 8C). Thus, the direct effect of *TENT4A* on *RAD18* mRNA appears to be dominant over the *PAXIP1* axis.

**Figure 8.**
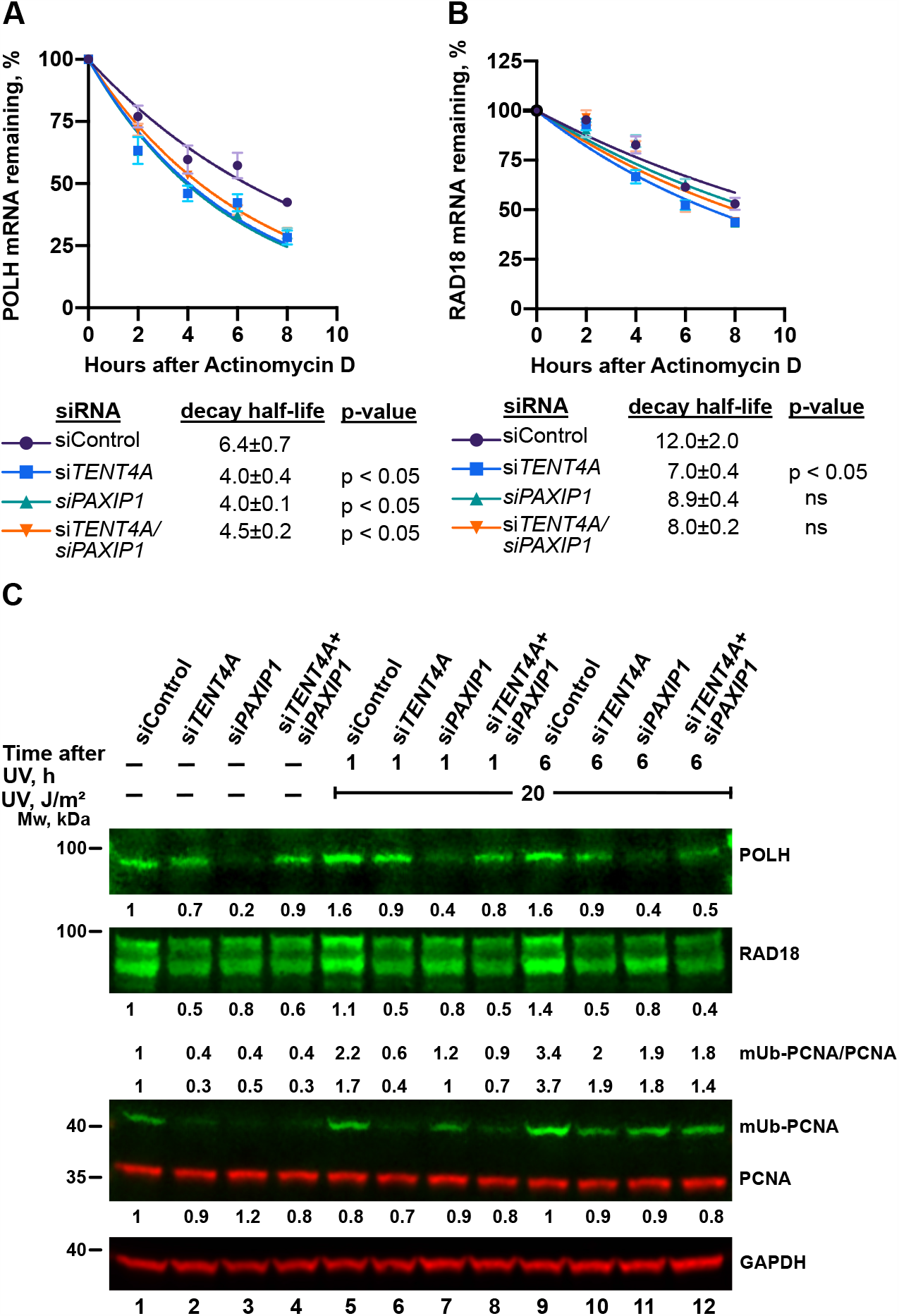
Effect of TENT4A and PAXIP1 on stability of TLS-related mRNAs and level of TLS-related proteins. (**A**) and (**B**) *TENT4A* and/or *PAXIP1* expression was knocked-down in MCF-7 cells for 48 h, after which the cells were treated with 5 μg/ml Actinomycin D for up to 8 h. The expression levels of *POLH and RAD18* were each determined by qPCR relative to the expression level at time 0. Half-life was calculated by using non-linear fit one phase decay curve equation and significance of the differences was calculated using Student’s *t*-test (one-tailed); Data are presented as mean ± SEM from three independent experiments. ns: *P*>0.05. (**C**) MCF-7 cells were transfected with *TENT4A* and/or PAXIP1 or non-targeting control siRNA for a final combined concentration of 100 nM. At 65 h post-transfection, the cells were UV-irradiated at 20 J/m^2^ and harvested 1 or 6 h post-irradiation. Whole cell extracts were fractionated by SDS–PAGE followed by western blot with indicated antibodies. Amounts are presented relative to those in extracts of unirradiated cells treated with siControl and the values are shown in the corresponding position of the blots.

### TENT4A and TLS genes are frequently mutated in endometrial cancer

The importance of TLS in tolerating DNA damage and affecting genetic stability prompted us to examine whether mutations in genes of the TLS pathway are over-represented in particular cancer types. To that end we used the TCGA database, and first analysed, for each cancer type, the percentage of the samples that contained mutations in at least one of the genes that we defined as a TLS-related genes group, containing *TENT4A, CYLD, NPM1, TENT4B, PAXIP1, PCNA, POLH, POLI, POLK, PRIMPOL, RAD18, REV1, REV3L* and *USP1*. As can be seen in Appendix Table S11, there is a big variation in the percent of samples with mutations in the TLS genes group, ranging from about 1-2% (e.g., Thyroid Carcinoma; THCA) up to 37% for Uterine Corpus Endometrial Carcinoma (UCEC; endometrial cancer). While as expected, cancer types with a higher overall number of mutations tend to exhibit a higher percentage of samples with mutations in TLS-related genes (e.g., Skin Cutaneous Melanoma, SKCM; Colon Adenocarcinoma, COAD; Lung Squamous Cell Carcinoma; LUSC; Appendix Table S11), endometrial cancer stands out with the highest occurrence of samples with mutations in TLS genes group, despite a higher median overall number of mutations in several other cancers (e.g., Bladder Cancer, BLCA; COAD; Lung Adenocarcinoma, LUAD; Appendix Table S11).

To probe the statistical significance of the high percentage of samples with mutations in the TLS genes group, we estimated the probability that this high percentage was obtained by chance. To that end we chose a control gene set of 14 random genes (the size of the TLS-related gene group), and examined in each cancer type the percentage of samples with mutations in at least one gene of this control gene set. This was repeated 1000 times and used to plot the chance distribution and calculate the probability of the TLS genes group (Appendix Fig S4, Appendix Table S11). Appendix Fig S4 shows also the average number of samples in each distribution, as well as the fraction of samples with TLS-related genes group in each cancer type. The fraction of the 1000 random sets which yielded a number of samples with mutations higher than the number of TLS mutations, represents the probability that the fraction of TLS-related genes group was obtained by chance, and was termed *P*-total value. Only three cancer types passed this statistical test with *P*-total<0.05, namely Acute Myeloid Leukemia (LAML), Thymoma (THYM) and UCEC (Appendix Table S11, Table 1A). For these three we tested the statistical significance of the difference between the number of samples with TLS mutations and the average number of samples with random mutation sets, using the Chi-squared test. Only LAML and UCEC passed this additional test with *P*<0.05 (Table 1A). Analysis of the fraction of samples with mutations in each TLS gene (Table 1B), shows that UCEC has indeed a considerable percentage of mutations in each of the TLS-related genes, including *TENT4A* (Table 1B), whereas in the LAML, the majority of samples contained mutations in a single gene, namely *NPM1* (Table 1B).

**Table 1.**
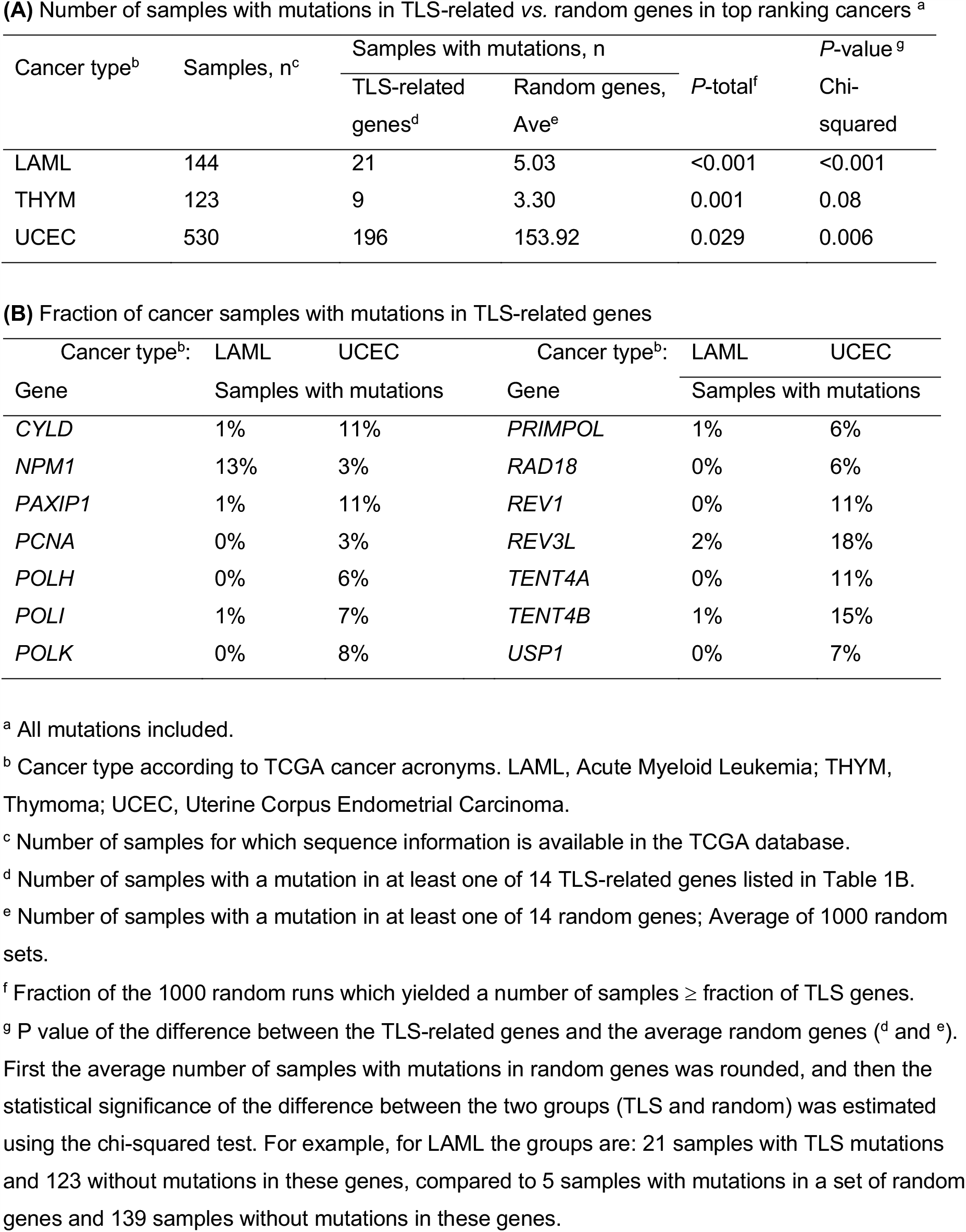
Analysis of the frequency of TLS-related gene mutations in cancer genomes ^a^.

## Discussion

DNA damage tolerance by TLS must be carefully regulated to balance between the beneficial effect of enabling overcoming replication obstacles, with minimal adverse effect of genetic stability. Indeed, no less than 17 genes were previously identified in our lab as new regulators of TLS (Ziv et al., 2014), amongst them *TENT4A* (*PAPD7*). While initially this gene was proposed to encode in yeast a DNA polymerase, it was subsequently shown to encode a poly(A) RNA polymerase in both yeast (Haracska et al., 2005) and human cells (Lim et al., 2018), with no detectable DNA polymerase activity, as confirmed in this study.

While little is known about the role of *TENT4A* in various biological processes, the current study shows that it has broad effects, and is involved in multiple pathways, most notably RNA processing and mRNA degradation, membrane-related pathways such as membrane budding, vesicle-mediated transport, vesicle organization etc., metabolism, proteolysis and cell division related processes (Fig 6). The finding that *TENT4A* is involved in TLS, which we has been studied in our lab over many years, prompted us to further explore the molecular mechanisms that underlie this involvement. Interestingly, the *TENT4A*-dependent regulation of TLS uncovered in the current study, spans several major components of the TLS machinery, i.e., RAD18, DTL, POLH and mUb-PCNA, in a complex interrelationship using both direct and indirect mechanisms (Fig 9). Interestingly, the *CYLD* tumor suppressor gene, previously identified as a TLS regulator (Ziv et al., 2014) turned out to be acting downstream to *TENT4A*, and directly regulated by it at the mRNA stability and translation levels, and *PAXIP1-AS2* was identified as a new significant *TENT4A*-regulated TLS regulator. Of note, *PAXIP1*, which *PAXIP1-AS2* regulates, is a paired box (PAX) gene with six BRCT domains, which is involved in development (Callen, Faryabi et al., 2012, Schwab, Smith et al., 2013) and responses to DNA damage (Mijic, Zellweger et al., 2017, Wang, Aroumougame et al., 2014). It is a suggested lung cancer tumor suppressor (Wu, Tian et al., 2018), and linked to breast and ovarian cancer, and response to chemotherapy (Jhuraney, Woods et al., 2016).

**Figure 9.**
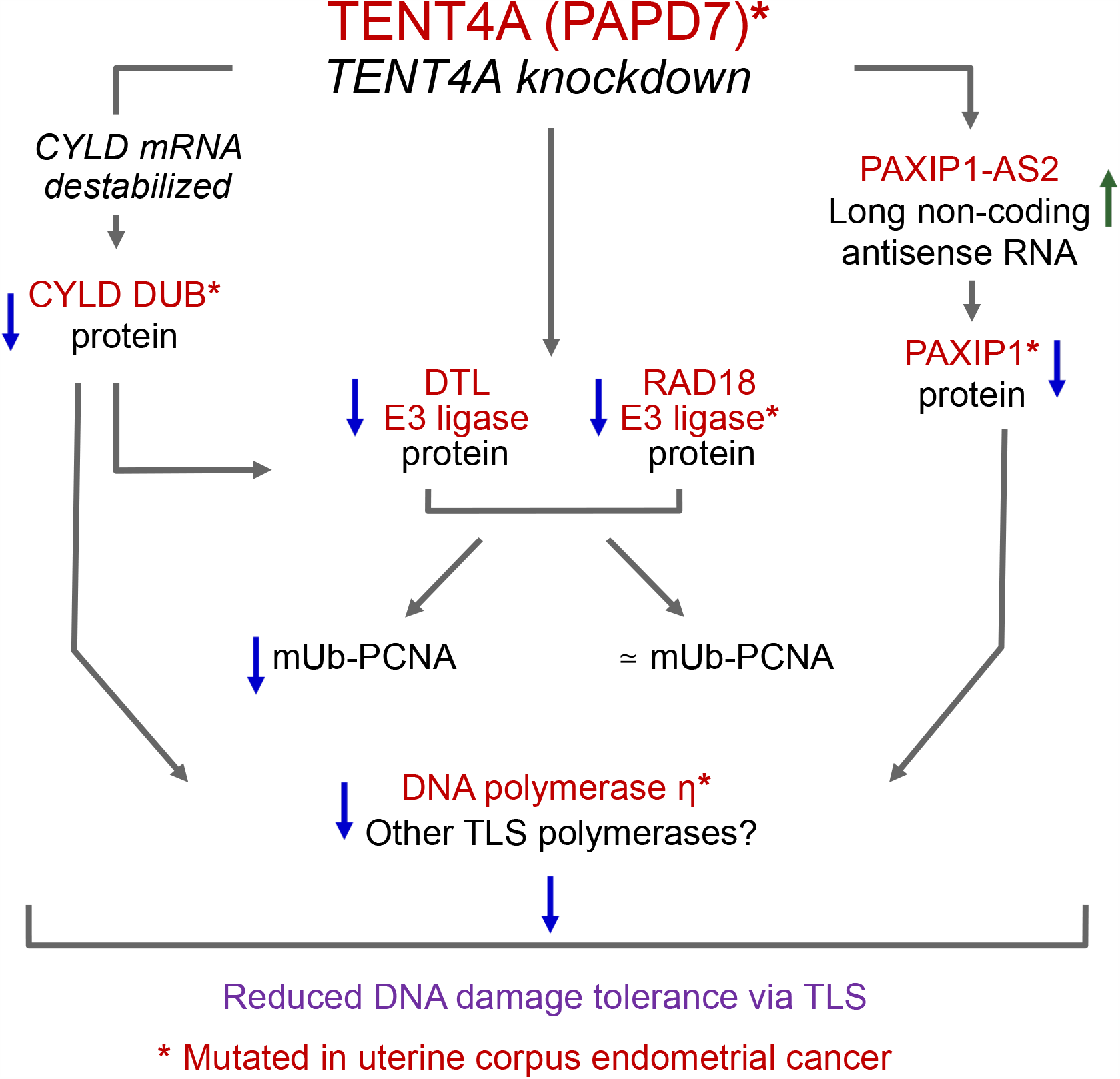
A scheme representing the three-branches of regulation of TLS by *TENT4A*. See text for details.

While *TENTA* directly regulates *RAD18* and *CYLD* via binding their mRNAs and controlling the length of their poly(A) tails and stability, consistent with its activity as non-canonical poly(A) RNA polymerase, *POLH* mRNA regulation appears to be indirect, since no binding of *POLH* mRNA to TENT4A protein was observed. It is likely mediated via the PAXIP1 protein (see below) which showed an effect similar and epistatic to *TENT4A*. Interestingly, *TENT4A* knockdown had also a global effect on translation, as indicated by the shift in ribosomes profile, and specifically manifested in the inhibited translation of the *POLH* and *CYLD* genes, but not *RAD18*.

Knocking down the expression of *TENT4A* or its downstream effector *CYLD* reduced the level of RAD18, and was therefore expected to be manifested in a decrease in mUb-PCNA. Unexpectedly, such a decrease was not necessarily observed, leading to the unravelling of an interesting relationship between TENT4A, RAD18 and DTL. First, in addition to RAD18, DTL is also regulated by *TENT4A*, whose knockdown strongly decreased the level of DTL protein. Second, knocking down the expression of *DTL*, caused a decrease in the amount of the RAD18 protein, in both unirradiated and UV-irradiated cells. This is not due to cross targeting of the siRNAs, since they are gene specific, raising the possibility that DTL acts upstream to RAD18. Because knocking down the expression of *CYLD* also reduced the amount of DTL, it is possible that the *CYLD* branch regulated by *TENT4A* operates in the order of *TENT4A* => *CYLD* => *DTL* => *RAD18*.

Interestingly, knocking down the expression of *RAD18* led to an *increase* in the amount of DTL protein, as if to compensate for the absence of this activity, possibly acting via a feedback loop. The fact that a reduction in the amount of RAD18 is not always manifested in a significant decrease in mUb-PCNA, may have been due to the remaining amounts of the RAD18, which were sufficient to carry out significant PCNA monoubiquitination, and/or there might exist a third, as yet unknown E3 ligase which monoubiquitinates PCNA.

As expected from its sequence complementarity, we found that *PAXIP1-AS2* is indeed a negative regulator of *PAXIP1* (Fig 7E). However, somewhat unexpectedly, this regulatory effect was not mediated via an effect on the amount or stability of *PAXIP1* mRNA (Fig 7C and D). It was previously reported that antisense long non-coding RNA can control translation (Carrieri, Cimatti et al., 2012), and a similar effect may explain the effect of *PAXIP1-AS2*.

*PAXIP1-AS2* overexpression, *PAXIP1* or CYLD knockdown each reduced the levels of POLH, and CYLD knockdown also reduced the level of RAD18, causing a significant decrease in TLS, despite little change in the level of mUb-PCNA. This decreased TLS was likely caused by the decreased level of POLH, which performs actual TLS reactions, with possible participation of RAD18, which in addition to the monoubiquitination of PCNA has a non-catalytic function in TLS, involving an interaction with POLH and guiding it to replication stalling sites (Durando, Tateishi et al., 2013, Huang, Zhou et al., 2018, Watanabe et al., 2004). The phenomenon, in which TLS was reduced but mUb-PCNA remained essentially unchanged, indicates that the formation of mUb-PCNA cannot be used as a surrogate for TLS, as is sometimes done, and may lead to erroneous conclusions.

TLS is involved in a variety of biological processes, most notably DNA damage tolerance (Livneh et al., 2016) and the generation of somatic hypermutation in the immune system (Casali, Pal et al., 2006). Dysregulation of TLS can cause an altered load of point mutations, or if TLS is inhibited it can lead to an increase in chromosomal aberrations due to the activity of alternative recombinational and repair events (Wittschieben, Reshmi et al., 2006). Of the 33 cancer types examined for occurrence of mutations in TLS-related genes, endometrial cancer is the only one which exhibits TLS mutations in a considerable fraction of samples, spread over all 14 genes, including *TENT4A* (Table 1), suggest that dysregulated or downregulated TLS is involved in endometrial carcinogenesis (Cancer Genome Atlas Research, Kandoth et al., 2013, Suhaimi, Ab Mutalib et al., 2016). It is yet unclear why this should be specific to endometrial cancer, however, dysregulation or inactivation of discrete DNA repair mechanisms is generally cancer type-specific, for reasons which are not fully understood (Akbari et al., 2015, Errol et al., 2006, Hanahan & Weinberg, 2011, Li et al., 2004, Marteijn et al., 2014, Modrich, 1994). Of note, the loss of TLS might sensitize cancer cells to DNA damaging chemotherapy, providing potential prognostic and therapeutic value (Gallo & Brown, 2019, Zafar & Eoff, 2017). To the best of our knowledge, this is the first report on such extensive somatic mutations in genes of the TLS pathway in sporadic cancers.

## Materials and Methods

### Cell cultures and transfections

Osteosarcoma U2OS, breast cancer MCF-7, human embryonic kidney HEK 293T and 293FT epithelial cells were cultured in DMEM (Gibco) supplemented with 2 mM L-Alanyl-L-Glutamine, 100 units/ml of penicillin, 100 µg/ml of streptomycin, 1 mM sodium pyruvate (Biological Industries), and 10% fetal bovine serum (HyClone). XPA (XP12RO) human fibroblasts derived from xeroderma pigmentosum patients were a gift from A. R. Lehmann (University of Sussex, Brighton, U.K.). XP12RO cells were cultured in MEM Eagle (Biological Industries) supplemented with 2 mM L-Alanyl-L-Glutamine, 100 units/ml of penicillin, 100 µg/ml of streptomycin (Biological Industries), and 15% fetal bovine serum (HyClone). The cells were incubated at 37°C in a humidified incubator with 5% atmosphere and periodically examined for mycoplasma contaminations by EZ-PCR test kit (Biological Industries). For plasmid transfection, cells were transfected with Lipofectamine 2000 (Invitrogen) or JetPRIME reagent (Polyplus) according to the manufacturer’s instructions. For siRNA transfection, the siGENOME SMART pool siRNA oligonucleotides, siGENOME Non-Targeting Control siRNA #5 (D-001210-05) and ON-TARGET*plus* SMART pool siRNA oligonucleotides and ON-TARGET*plus* NON-targeting pool (D-001810) (Dharmacon) were transfected with 25 or 50 nM siRNA by Lipofectamine RNAimax (Invitrogen) for 48 to 72 h following the manufacturer’s protocol. For combinatorial knockdown, an equal amount of siRNAs against each gene was mixed to have a final combined concentration of 100 nM for the double knockdown and 150 nM for the triple knockdown, unless otherwise stated. siRNAs used in this study are listed in Appendix Table S12.

### Generation of lentiviral stable cell lines

Lentiviral pLKO.1-puro empty vector control plasmid and human *TENT4A* shRNA pLKO.1-puro plasmid (clone TRCN0000053036, target sequence CCAACAATCAGACCAGGTTTA) were obtained from Sigma (Mission shRNA library). Lentiviruses were produced in 293FT cells, by co-transfecting pLKO-derived plasmids and the second-generation packaging plasmids using JetPRIME. Lentiviral particles were harvested 48 h post-transfection. The resulting lentiviral particles were used to infect the U2OS cells. 48 h post-infection, 2 µg/ml of puromycin (InvivoGen) was added to select for infected cells. After two weeks of selection, individual colonies were isolated and tested for knockdown of *TENT4A* by quantitative real-time PCR (qPCR).

### TLS assay in cultured mammalian cells

The TLS assay was described earlier (Diamant, Hendel et al., 2012, Shachar et al., 2009, Ziv, Diamant et al., 2012). The TLS gap-plasmid transfection was performed by co-transfecting with a mixture containing 50 ng of a lesions plasmid (Kan^R^), 50 ng of a gapped plasmid without a lesion (Cm^R^), and 1900 ng of the carrier plasmid pUC18, using Lipofectamine 2000. The cells were incubated for 4 h (for the TTCPD gap-lesion plasmid), 20 h (TT 6-4PP), 16 h (BP-G), or 6 h (cisPt-GG) to allow TLS. The plasmids were extracted using alkaline lysis conditions followed by renaturation, such that only covalently closed plasmids remained nondenatured. A fraction of the purified DNA was used to transform a TLS-defective *E. coli recA* strain JM109, which was then plated on LB-kan and LB-cm plates. The efficiency of gap repair was calculated by dividing the number of transformants obtained from the gap-lesion plasmid (number of colonies on LB-kan plates) by the number of corresponding transformants obtained with the control gapped plasmid GP20-cm (number of colonies on LB-cm plates). To obtain precise TLS extents, the plasmid repair extents were multiplied by the fraction of TLS events out of all plasmid repair events, based on the DNA sequence analysis of the plasmids from Kan^R^ colonies. To determine the DNA sequence changes that have occurred during plasmid repair, sequence analysis was carried using the TempliPhi DNA Sequencing Template Amplification Kit and the BigDye Terminator v1.1 Cycle Sequencing Kit. Reactions were analyzed by capillary electrophoresis on an ABI 3130XL Genetic Analyzer from Applied Biosystems.

### RNA isolation and qPCR

Total RNA was isolated using RNeasy plus mini kit (Qiagen) according to the manufacturer’s instructions, including treatment with RNase-free DNase I (Qiagen). cDNA was synthesized using High Capacity cDNA Reverse Transcription Kit (Applied Biosystems). qPCR was performed using qPCRBIO SyGreen Blue Mix (PCR Biosystems) and run on a QuantStudio™ 6 Flex Real-Time PCR System (Applied Biosystems). Primer sequences used were predesigned KiCqStart SYBR® Green primers (Sigma-Aldrich) and are included in Appendix Table S13. The expression of the indicated transcript was normalized to endogenous reference control GAPDH according to ΔΔCt method using the DataAssist software v3.0 (Applied Biosystems).

### mRNA stability assay

The expression of *TENT4A, TENT4B, PAXIP1* or *TENT4*A plus *TENT4B*, or *TENT4A* plus *PAXIP1* was knocked-down for 48 h in MCF-7 cells. To measure mRNA stability, 5 µg/ml Actinomycin D (Sigma-Aldrich) was added to the growth medium to inhibit transcription, cells were harvested at the indicated time points and mRNA expression was measured by qPCR. The half-life of the mRNA was calculated by the non-linear fit, one phase decay curve equation using GraphPad Prism 8 software.

### RNA-Immunoprecipitation (RNA-IP) and extension poly(A) test (ePAT)

RNA-IP to assay TENT4A/RNA interactions was performed as described (Keene, Komisarow et al., 2006), and the ePAT assay was performed as described with some modifications (Janicke et al., 2012), and the two are presented in the Appendix Materials and Methods and Appendix Tables S13 and S14.

### UV-irradiation, Whole cell extracts, and Western blotting

When indicated, cells were rinsed in Hanks’ buffer and irradiated in Hanks’ buffer with UV-C using a low-pressure mercury lamp (TUV 15W G15T8, Philips) at a dose rate of 0.2 J/m^2^/s. UV dose rate was measured using an UVX Radiometer (UVP) equipped with a 254-nm detector. After irradiation, Hanks’ buffer was removed and the cells were incubated in fresh growth medium for additional time before harvest.

Cells were resuspended in CelLytic M lysis buffer (Sigma-Aldrich) containing protease inhibitor cocktail, phosphatase inhibitor 2 and 3 cocktail (Sigma-Aldrich), 2.5 mM MgCl_2_, and 50 units/ml Benzonase (Merck), incubated on ice for 30 min followed by centrifugation. The supernatant was collected and the protein concentration was determined using the Bradford protein assay (Bio-Rad). The extracts were fractionated on a 4-20% ExpressPlus™ PAGE gel (Genscript) using SDS-MOPS buffer and transferred onto nitrocellulose membrane (BioTrace™ NT Nitrocellulose transfer membrane, Pall Corporation), followed by blocking of membranes with Odyssey Blocking buffer (LI-COR Biosciences, Lincoln, NE, USA) for 1 h at room temperature. The membranes were incubated with primary antibodies diluted in Odyssey Blocking Buffer–0.1% Tween-20 overnight at 4 °C. The membranes were washed 3×10 min with Tris-buffered saline with 0.1 % Tween-20 (TBST) and further incubated for 1 h at room temperature in the IRDYE-680 or 800 conjugated secondary antibodies diluted in Odyssey Blocking Buffer–0.1% Tween-20. Membranes were washed 3×10 min TBST and finally rinse with TBS to remove residual Tween-20. The membranes were imaged on a LI-COR Odyssey Fc Imager (LI-COR Biosciences) and the bands were quantified by ImageStudio v 5.2.5 (LI-COR Biosciences) software.

### Antibodies

Commercially available antibodies used were as follows: rabbit anti-RAD18 (Cell Signaling, 9040S, dilution 1:2000); rabbit anti-ubiquityl-PCNA (Lys164) (Cell Signaling, 13439S, dilution 1:2000); rabbit anti-GAPDH (Cell Signaling, 5174S, dilution 1:10000); mouse anti-GAPDH (Milipore, MAB374, dilution 1:10000); rabbit anti-POLH (Cell Signaling, 13848S, dilution 1:1000); rabbit anti-CYLD (Cell Signaling, 8462S, dilution 1:1000); mouse anti-PCNA (PC-10) (Santa Cruz, sc-56, dilution 1:1000); mouse anti-FLAG M2 (Sigma-Aldrich, F1804, dilution 1:1000); rabbit anti-PTIP (PAXIP1) (Bethyl, A300-370A, dilution 1:2000); rabbit anti-DTL (Bethyl, A300-948A, dilution 1:1000); rabbit anti-TRF4 (H-172) (TENT4A) (Santa Cruz, sc-98490, dilution 1:500); anti-MBP monoclonal antibody, HRP conjugated (New England Biolabs, E8038S, dilution 1:10000); IRDye 680RD goat anti-mouse IgG (LI-COR, 926-68070, dilution 1:10000); IRDye 800CW goat anti-mouse IgG (LI-COR, 926-32210, dilution 1:10000); IRDye 680RD goat anti-rabbit IgG (LI-COR, 926-68071, dilution 1:10000); IRDye 800CW goat anti-rabbit IgG (LI-COR, 926-32211, dilution 1:10000).

### RNA-seq and Bioinformatics

The sources of RNA were U2OS cells in which the expression of *TENT4A* was knocked-down using lentivirus-mediated shRNA combined with siRNA against *TENT4A*, and human XP12RO cells transfected with 25 nM of si*TENT4A* and siControl by Lipofectamine RNAimax for 48 h. RNA was extracted from three biological replicates, the quality assessed on the 2200 TapeStation system (Agilent). Library preparation and sequencing were performed at the Crown Genomics Institute of the Nancy and Stephen Grand Israel National Center for Personalized Medicine at the Weizmann Institute of Science. Briefly, sequencing libraries were prepared using TruSeq® Stranded Total RNA (Illumina) (Cat # RS122-2301, RS-122-2302) for ribosomal-depletion libraries. After rRNA depletion from 500 ng total RNA with rRNA Ribo-Zero Gold removal mix, cDNA was performed followed by second-strand synthesis with dUTP instead of dTTP. Then, A base addition, adapter ligation, UDG treatment and PCR amplification steps were performed. Sequencing was performed on Illumina HiSeq2500 machine. The median sequencing depth was 45 million reads. Poly-A/T stretches and Illumina adapters were trimmed from the reads using cutadapt; resulting reads shorter than 30bp were discarded. Reads were mapped to the Homo Sapiens GRCh38 reference genome using STAR (version 2.4.2a), supplied with gene annotations downloaded from Ensembl (release 88).The alignEndsType was set to EndToEnd and outFilterMismatchNoverLmax was set to 0.04. Expression levels for each gene were quantified using htseq-count (the stranded option was set to reverse), using the gtf above. Differential expression analysis was performed using DESeq2 (version 1.10.1). The betaPrior, Cook’s distance cutoff and independent filtering parameters were set to False. Raw P values were adjusted for multiple testing using the procedure of Benjamini and Hochberg (Benjamini & Hochberg, 1995). We defined significantly differential expressed genes in U2OS and XP12RO cell as those with FDR ≤ 0.05, absolute fold change ≥ 2 and a count of at least 30 at least in one of the samples.

### Gene set enrichment analysis (GSEA)

We performed GSEA using pre-ranked algorithm with rank determined by −log_10_(FDR q-value) * sign (fold change). The Java GSEA Desktop Application version 3 from the Broad Institute was used and the enrichment statistic parameter set to classic but other parameters remained at their default values. Enrichment analysis was performed and leading edge gene sets were determined by using GO processes (c5.bp.v6.1.symbols.gmt) from Molecular Signature Database (MSigDB) v6.1. Gene sets were considered significantly enriched at FDR<0.05. We further performed enrichment analysis in Pathway Studio MammalPlus V 12 (Elsevier) by GSEA and Fisher’s Exact Test for genes of U2OS and XP12RO.

### TCGA database mutation analysis

The R package ‘TCGAbiolinks’ (Colaprico, Silva et al., 2016) was used to download all the MAF (Mutation Annotation Format) files related to the 33 cancer types from TCGA database. Loading and summarizing the MAF files for each cancer was done using the R package ‘maftools’ (Mayakonda, Lin et al., 2018). For the analysis of each cancer type, MAF files from all four pipeline analysis available in the TCGA database (“muse”, “varscan2”, “somaticsniper”, “mutect2”) were merged using the ‘merge_mafs’ function from ‘maftools’. The variant classifications, “Frame_Shift_Del”, “Frame_Shift_Ins”, “Splice_Site”, “Translation_Start_Site”, “Nonsense_Mutation”, “Nonstop_Mutation”, In_Frame_Del”, “In_Frame_Ins”, “Missense_Mutation”,”3’UTR”, “5’UTR”, “3’Flank”, “Targeted_Region”, “Silent”, “Intron”,”RNA”, “IGR”, “Splice_Region”, “5’Flank”, “lincRNA”, “De_novo_Start_InFrame”, “De_novo_Start_OutOfFrame”, “Start_Codon_Ins”, “Start_Codon_SNP”, “Stop_Codon_Del”, were used in the analysis of mutations. Mutations frequencies were calculated as the number of mutated samples divided by the total number of samples for each gene per cancer.

### Statistical analysis

The statistical results were obtained from at least three independent biological replicates unless otherwise stated. All results were presented as mean ± SEM. *P* values were obtained via the Student’s t-test (two-tailed), unless otherwise stated, using GraphPad Prism 8.0 software. *\*P* < 0.05, *\*\*P* < 0.01, *\*\*\*P* < 0.001.

See Appendix for additional Materials and Methods.

## Supporting information

Supplemental RNA-seq data 1

Supplemental RNA-seq data 2

## Data availability

The RNA-seq data discussed in this publication have been deposited in NCBI’s Gene Expression Omnibus (Edgar, Domrachev et al., 2002) and are accessible through GEO Series accession number GSE141870 (https://www.ncbi.nlm.nih.gov/geo/query/acc.cgi?acc=GSE141870).

## Acknowledgements

This work was supported by the Flight Attendant Medical Research Institute, Florida, USA (FAMRI #032001 to Z.L. and T.P.E.); the Israel Science Foundation [#684/12 to Z.L.) and the Minerva Foundation (#120855) with funding from the Federal German Ministry for Education and Research (to Z.L.). Funding for open access charge: Flight Attendant Medical Research Institute, Florida, USA. Z.L. is the incumbent of the Maxwell Ellis Chair for Biomedical Research. R.D. is the incumbent of the Ruth and Leonard Simon Professorial Chair of Cancer Research. G.F. is the Incumbent of the David and Stacey Cynamon Research fellow Chair in Genetics and Personalized Medicine.

## Author contribution

UmSw designed, performed and analysed most experiments, and participated in writing the manuscript. GF performed the RNAseq analysis and commented on the manuscript. UrSe performed polysome profiling experiments, analysed their data, and commented on the manuscript. ASP and RR performed the mutational analysis based on the TCGA data base. CE, TC and NS provided oligonucleotides and commented on the manuscript. TPE, participated in designing experiments, analysis of the data, and writing the manuscript. RD provided advice, contributed to analysis of the results and commented on the manuscript. ZL conceived, devised and supervised the study, and wrote the manuscript.

