## Supplemental Methods, tables and figures for "TENT4A non-canonical poly(A) polymerase regulates DNA-damage tolerance via multiple pathways which are mutated in endometrial cancer"

**Table of Contents** **Pages**

| Table of Contents | Pages |
|---|---|
| Appendix Materials and methods | 2-7 |
| Appendix Table S1 | 8 |
| Appendix Table S2 | 9 |
| Appendix Table S3 | 10 |
| Appendix Table S4 | 11 |
| Appendix Table S5 | 12 |
| Appendix Table S6 | 13 |
| Appendix Table S7 | 14 |
| Appendix Table S8 | 15 |
| Appendix Table S9 | 16 |
| Appendix Table S10 | 17 |
| Appendix Table S11 | 18-19 |
| Appendix Table S12 | 20 |

Appendix Table S13 21

Appendix Table S14 22

Appendix Figure S1 23

Appendix Figure S2 24

Appendix Figure S3 25

Appendix Figure S4 26

References 27


**Appendix Materials and Methods**

**Plasmid construction.** *TENT4A* expression plasmid. The MGC human *TENT4A* sequence-verified cDNA (Accession: BC117137, Clone ID: 40125688) containing pCR4-TOPO was purchased from Open Biosystem. Extended N-terminal sequences of TENT4A (230 aa) was codon-optimized and synthesized as gBlock gene fragment (Integrated DNA technologies). Then, the extended sequences were subcloned into original plasmids of *TENT4A* (542 aa) by using Gibson assembly (New England Biolabs). The coding sequence of *TENT4A* was amplified and cloned into modified mammalian expression vector pLEXm vector (gift from Wei Yang, National Institutes of Health, Bethesda, USA) downstream of the chicken-beta actin promoter and Kozak sequence containing 8×His tag (His8) and maltose-binding protein (MBP) in tandem by Gibson assembly. D277A and D279A mutations in TENT4A were introduced by Q5® Site-Directed Mutagenesis Kit (New England Biolabs) to produce catalytically dead TENT4A. The mutagenic primers for D277A and D279A mutations in TENT4A were designed by NEBaseChanger (New England Biolabs). Similarly, C' FLAG-TENT4A was constructed by inserting FLAG tag in the C' terminus TENT4A by PCR and the C' FLAG TENT4A fragment was transferred to pcDNA3 vector by Gibson assembly.

*PAXIP1-AS2* expression plasmid. Homo sapiens *PAXIP1* antisense RNA 2 (*PAXIP1-AS2*), transcript variant 1, long non-coding RNA sequence was obtained from the University of

California Santa Cruz (UCSC) genome browser. The *PAXIP1-AS2* gencode transcript (ENST00000449486.1) consists of 2121 bp. The MGC human *PAXIP1-AS2* sequence-verified cDNA (Accession: BC033053, Clone ID: 5259766) containing pBluescriptR purchased from Open Biosystem. 1-1037 bp of *PAXIP1-AS2* transcript was PCR amplified from cDNA and the remaining 1048 bp was synthesized as gBlock gene fragment (Integrated DNA technologies). These PCR products and a linearized pCMV SPORT6 vector were assembled by Gibson assembly reaction to create a PAXIP1-AS2 pCMV SPORT6 expression plasmid. All the plasmid maps construction and Gibson assembly simulation were performed by SnapGene v3.3.1 software (GSL Biotech LLC).

**Protein expression and purification.** HEK 293T cells at approximately 70-80% confluence in T175 culture flasks were transfected with 25 μg of 8xHis-MBP TENT4A pLEXm and 8xHis-MBP TENT4A D277A, D279A pLEXm plasmids using 50 μl of JetPRIME reagent according to the manufacturer's instructions. Thirty-six hours after transfection cells were washed with cold PBS, harvested, centrifuged, and the pellet was snap-freeze in liquid nitrogen and stored in a deep-freezer at -80°C. Cell pellet (~6 g) was resuspended in 25 ml of CelLytic M lysis buffer (Sigma-Aldrich) containing 300 mM NaCl, Protease inhibitor cocktail (1:100, Sigma), 5 mM β-mercaptoethanol, 2.5 mM $MgCl_2$, and 50 units/ml Benzonase (Merck), and the tube was rotated for 1 h at 4°C. After centrifuging at 40,000 g in a Sorval RC-6 centrifuge (Thermo) for 1 h, the lysates were adjusted to binding buffer conditions, namely 20 mM Tris-Cl (pH 7.5), 300 mM NaCl, 5% (vol/vol) glycerol, 0.01% IGEPAL CA-630 (Sigma), 5 mM β-mercaptoethanol and 5 mM Imidazole. The lysates were then mixed with 2 ml of Co-NTA resin (Cube Biotech, Germany), which were preequilibrated with the binding buffer, and incubated with rotation for 1 h at 4°C. After pouring the resin into a column and washing thoroughly with 50 column volume of wash buffer (20 mM Tris-Cl (pH 7.5), 300 mM NaCl, 5% (vol/vol) glycerol, 0.01% IGEPAL CA-630, 5 mM β-mercaptoethanol and 10 mM Imidazole), the proteins were eluted with 5 column volume of elution buffer (20 mM Tris-Cl (pH 7.5), 300 mM NaCl, 0.01% IGEPAL CA-630, 250 mM imidazole). The elutes were buffer exchanged with exchange buffer (20 mM Tris-Cl (pH 7.5), 200 mM NaCl, 1 mM EDTA, 5% (vol/vol) glycerol, 0.01% IGEPAL CA-630, 0.5 mM Tris(2-carboxy- ethyl)phosphine (TCEP)) by PD-10 desalting column (GE Healthcare), and then NaCl concentration was increased to 1 M. The elutes were mixed with 1 ml of preequilibrated amylose resin (NEB), and incubated with rotation for 2 h at 4°C. After pouring the resin into a column and thoroughly washing with 200 column volume of MBP wash buffer (20 mM Tris-Cl (pH 7.5), 1 M NaCl, 1 mM EDTA, 5% (vol/vol) glycerol, 0.01% IGEPAL CA-630, 0.5 mM TCEP), the proteins were eluted with 5 column volume of MBP elution buffer (20 mM Tris-Cl (pH 7.5), 150 mM NaCl, 0.5 mM EDTA, 5% (vol/vol) glycerol, 0.01% IGEPAL CA-630, 1 mM TCEP and 20 mM Maltose). The final elutes were concentrated with Vivaspin Turbo 4, 50K MWCP PES (Sartorius), aliquoted, snap-frozen in liquid nitrogen, and stored at -80°C.

**Poly(A) RNA polymerase assays.** Poly(A) RNA polymerase assays were performed in 10 µl reaction buffer containing 25 mM Tris-Cl (pH 8), 20 mM KCl, 2 mM DTT, 0.02% NP-40, 2% glycerol, 50 µg/ml bovine serum albumin, 1 mM of each ATP, GTP, CTP, and UTP, 1 mM $MnCl_2$ or 5 mM $MgCl_2$, 1 pmol of $^{32}P$-labeled synthetic RNA oligonucleotide and 5-40 ng of partially purified TENT4A or Mutant protein. Reactions were incubated at 37°C for 20 min and stopped by the addition equal volume of a mixture of loading buffer containing 25 mM EDTA, 95% formamide, 0.25% bromophenol blue, and 0.25% xylene cyanol. For positive and negative controls, parallel reactions were also carried out with *E .coli* poly(A) polymerase (*E.coli* PAP) and no added proteins. The reaction products were resolved on 15% polyacrylamide gels containing 8 M urea, after which they were dried and visualized using a Personal Molecular Imager System (PMI) and Image Lab software (Bio-Rad).

**DNA polymerase assays.** DNA polymerase assays were performed in 10 µl reaction buffer containing 25 mM Tris-Cl (pH 8), 20 mM KCl, 2 mM DTT, 0.02% NP-40, 2% glycerol, 50 µg/ml bovine serum albumin, 100 µM of each dATP, dGTP, dCTP, and dTTP, 1 mM $MnCl_2$ or 5 mM

$MgCl_2$, 0.5 pmol of a $^{32}P$-labeled 15-nt primer annealed to a 60-nt template, and 5-40 ng of partially purified TENT4A or Mutant protein. Assay mixtures were assembled on ice, then incubated at 30°C for 20 min, and stopped by the addition equal volume of a mixture of loading buffer containing 25 mM EDTA, 95% formamide, 0.25% bromophenol blue. Analysis of the reaction was performed as described above for the poly(A) RNA polymerase assay.

**RNA-Immunoprecipitation (RNA-IP).** RNA-IP to assay TENT4A/RNA interactions were performed as described (Keene *et al*, 2006). HEK 293T cells were transfected with FLAG-TENT4A pcDNA3 and pcDNA3 empty vector by using JetPRIME transfection reagent according to the manufacturer's instruction. After 24 h of transfections, cells were rinsed with ice-cold 1×PBS, scrapped and lysed in a pellet-equal volume of ice-cold polysome lysis buffer (10 mM HEPES-KOH (pH 7.0), 100 mM KCl, 5 mM $MgCl_2$, 0.5% IGEPAL CA-630 supplemented by 1 mM DTT, Protease inhibitor cocktail, 0.4 mM Vanadyl ribonucleoside complexes (NEB) and 100 units/ml RNaseOUT (Invitrogen)), and stored at -80°C. Prewashed Dynabeads-Protein G (50 µl per reaction tube) resuspended in 50 mM Tris-Cl (pH 7.4), 150 mM NaCl, 1 mM $MgCl_2$, and 0.05% IGEPAL CA-630 (NT2 buffer) supplemented with 100 units/ml RNAseOUT were incubated with monoclonal anti-FLAG M2 antibody (Sigma-Aldrich) with gentle rotation for 1 h at room temperature. Subsequently, the magnetic beads were washed five times with NT2 buffer (1 ml each) and incubated with cell lysate supplemented with 200 units RNaseOUT, 1 mM DTT, and 20 mM EDTA (pH 8.0) in NT2 buffer for 3 h at 4°C tumbling end over end. Magnetic beads were then washed three times with ice-cold NT2 buffer, then twice with 300 mM NaCl containing NT2 buffer, and finally with NT2 buffer. FLAG-TENT4A/RNA complexes were eluted with excess FLAG peptide (Sigma-Aldrich) and bound RNA was isolated with TRI (Sigma-Aldrich) and purified according to the manufacturer's instructions. The amount of RNA was then quantified using NanoDrop (Thermo Fisher Scientific), and cDNAs were synthesized using the qScript cDNA Synthesis Kit for RT-PCR (QuantaBio). Equal amounts of cDNA were subjected to qPCR using KAPA SYBR® FAST qPCR Master Mix (2X) (KapaBiosystems) using the gene specific pair of primers as was described in Appendix Table S13.

**Extension Poly(A) Test (ePAT) assay.** ePAT was performed as described with some modifications (Janicke *et al*, 2012). Briefly, 1 µl of 100 µM DNA oligonucleotide (anchor primer) was annealed to 1 µg RNA (extracted from *TENT4A* or *TENT4B* knocked-down MCF-7 cells) by incubating at 80°C for 5 min then cooling down to room temperature. A master mix containing 1 X SuperScript III buffer, 5 mM DTT, 0.5 mM dNTPs, 40 units RNase Inhibitor (NEB) and 5 units of Klenow polymerase (NEB) was added to give a final reaction volume of 20 µl, mixed and incubated at for 1 h at 37°C, followed by 5 min at 80°C to inactivate the polymerase and cooling to 55°C for 1 min. The tubes were maintained at 55°C while 200 units of SuperScript III Reverse Transcriptase (Invitrogen) was added. The samples were mixed and incubated for 1 h at 55°C followed by 5 min at 80°C to inactivate the reverse transcriptase. A TVN-PAT cDNA control was generated by the same method using the TVN-PAT primer, which contains two 3′ variable bases, V and N, which lock the primer to the polyadenylation site during the reverse transcription step, which was performed at 42°C for 1 h and 52°C for 1 h. PCR amplification of the ePAT and TVN-PAT cDNA was performed in a 20 µl reaction containing 1 µl of cDNA, 1X AmpliTaq Gold 360 master mix (Applied Biosystems), 0.2 µM forward primer (gene specific primer), and 0.2 µM reverse primer (anchor primer). Reactions were incubated at 95°C for 10 min then 35 cycles of 95°C for 30 sec, 60°C for 30 sec, 72°C for 30 sec, and finally 7 min at 72°C. PCR amplicons were visualized by loading 10 µl of each reaction on to a 2% high-resolution agarose gel (Ultra-pure 1000; Invitrogen) containing SYBR Safe (Invitrogen) in Li-COR Odyssey Fc Imaging System (LI-COR Biosciences, Lincoln, NE, USA). To estimate the PCR product sizes and to quantify the intensity mass of PCR amplicons from such gels, the band intensity and migration was determined relative to a 100-bp ladder (SMOBIO Technologies) using an Image J software with line-width plugin and GraphPad Prism V.8. The primers used in this study are presented in Appendix Table S14.

**Polysomes Profiling**

MCF-7 cells were transfected with 50 nM siRNA against *TENT4A* or siGENOME Non-Targeting Control siRNA #5. Seventy-two hours later the cells were incubated with 100 µg/ml cycloheximide for 5 min and then washed twice with cold polysome buffer containing 20 mM Tris-Cl (pH 8), 140 mM KCl, 5 mM $MgCl_2$, and 100 µg/ml cycloheximide. The cells were harvested and lysed with 500 µl polysome buffer with 0.5% Triton X-100, 0.5% sodium deoxycholate, 1.5 mM DTT, 150 units RNAse inhibitor, and 5 µl of protease inhibitor cocktail. The lysed samples were centrifuged at 12,000g at 4°C for 5 min. The cleared lysates were loaded onto 10-50% sucrose gradient and centrifuged at 288,000 g in a SW41 rotor for 90 min at 4°C. Gradients were fractionated with the optical density at 254nm being continuously recorded using ISCO absorbance detector UA-6. RNA was isolated using Trizol and DirectZol RNA mini-prep kits (Zymo Research). cDNA was prepared using High-capacity cDNA reverse transcription kit (Applied Biosystems). qPCR was performed with qPCRBIO SyGreen Blue Mix in Quant 6 real PCR system and the % mRNA present in each fraction was calculated using the 2^-Ct method.

**Appendix Table S1. TLS across a site-specific TT 6-4PP or BP-G in *TENT4A* knocked-down U2OS cells.**

| Sample | Lesion | Transformants [a] | | Plasmid repair [b], % | TLS [c], % | Relative TLS,% |
|---|---|---|---|---|---|---|
| | | $Kan^R$ | $Cm^R$ | | | |
| sh/siControl | TT 6-4PP | 85 | 665 | 12.1±0.4 | 10±0.3 | 100±3 |
| sh/si*TENT4A* | | 28 | 415 | 6.7±0.8 | 5.2±0.6 | 52±6 |
| sh/siControl | BP-G | 223 | 1595 | 13.7±0.9 | 12.6±0.8 | 100±7 |
| sh/si*TENT4A* | | 58 | 894 | 6.1±0.2 | 5.4±0.2 | 43±2 |

[a] Results of a typical experiment are presented

[b] Plasmid repair is defined as the number of $Kan^R$ colonies divided by the number of $Cm^R$ colonies. The average of three experiments are presented.

[c] TLS values are obtained by multiplying plasmid repair values by the fraction of TLS events, as deduced from the DNA sequence analysis.

**Appendix Table S2. Analysis of mutations formed during TLS across a TT 6-4 PP in *TENT4A* knocked-down U2OS cells.**

| Event type | Sequence opposite TT 6-4PP in 5’-AA*TT*GT-3’ | sh/siControl | sh/si*TENT4A* |
|---|---|---|---|
| Accurate TLS | 3'-TTAACA-5' | 57 | 52 |
| Mutagenic TLS | | | |
| Misinsertion opposite 3' of lesion | 3'-TTA**G**CA-5' | 4 | 5 |
| | 3'-TTA**T**CA-5' | 5 | 7 |
| | 3'-TTA**C**CA-5' | 3 | 1 |
| Double misinsertions | 3'-T**A**A**G**CA-5' | 1 | 0 |
| | 3'-**C**TA**G**CA-5' | 0 | 1 |
| | 3'-**C**TA**T**CA-5' | 0 | 2 |
| | 3'-**C**TA**C**CA-5' | 0 | 1 |
| Misinsertion opposite 5' of lesion | 3'-TT**G**ACA-5' | 3 | 1 |
| | 3'-TT**T**ACA-5' | 2 | 0 |
| Tandem-double opposite lesion | 3'-TT**TG**CA-5' | 1 | 0 |
| Non-TLS events | | 16 | 20 |
| Total clones analyzed | | 92 | 90 |

Plasmids were extracted from $Kan^R$ colonies obtained in the experiments described in methods, and subjected to DNA sequence analysis. The 6-4 PP lesion is shown in italics (*TT*). Insertion of incorrect nucleotides is indicated by bold type letters. The two nucleotides incorporated opposite to the site of the TT 6-4 PP are underlined.

Non-TLS events include big deletions and insertion of sequences from the control plasmid.

**Appendix Table S3. Analysis of mutations formed during TLS across a BP-G in *TENT4A* knocked-down U2OS cells.**

| Event type | Sequence opposite 5'-GTTCGT*G*ACGTG-3' | Number of events | |
|---|---|---|---|
| | | sh/siControl | sh/si*TENT4A* |
| Accurate TLS | 3'-ACT-5' | 65 | 66 |
| Mutagenic TLS | | | |
| | 3'-A**A**T-5' | 9 | 6 |
| | 3'-A**T**T-5' | 2 | 2 |
| | 3'-A**G**T-5' | 4 | 7 |
| | 3'-AC**C-5'** | 3 | 1 |
| Non-TLS events | | 7 | 10 |
| Total clones analyzed | | 90 | 92 |

Plasmids were extracted from $Kan^R$ colonies obtained in the experiments described in methods, and subjected to DNA sequence analysis. The BP-G lesion is shown in italics (*G*). Insertion of an incorrect nucleotide is indicated by a bold type letter. The nucleotide incorporated opposite to the site of the BP-G is underlined.

Non-TLS events include big deletions and insertion of sequences from the control plasmid.

**Appendix Table S4. TLS across a site-specific cisPt-GG or TT-CPD lesion in *TENT4A* and/or *TENT4B* knocked-down MCF-7 cells.**

| Sample | Lesion | Transformants [a] | | Plasmid repair [b], % | TLS [c], % | Relative TLS,% |
|---|---|---|---|---|---|---|
| | | $Kan^R$ | $Cm^R$ | | | |
| siControl | cisPt-GG | 102 | 217 | 45±2 | 44±2 | 100±4 |
| si*TENT4A* | cisPt-GG | 19 | 79 | 25±1.5 | 23±2 | 52±3 |
| si*TENT4B* | cisPt-GG | 65 | 174 | 39±3.6 | 38±8 | 87±8 |
| si*TENT4A/4B* | cisPt-GG | 42 | 170 | 28±2.7 | 26.4±2 | 60±5 |
| siControl | TT-CPD | 91 | 136 | 71±5 | 65±5 | 100±7 |
| si*TENT4A* | TT-CPD | 32 | 88 | 34±4 | 31±4 | 48±5 |
| si*TENT4B* | TT-CPD | 105 | 160 | 64±5 | nd | nd |
| si*TENT4A/4B* | TT-CPD | 68 | 143 | 48±2 | nd | nd |

[a] The results of a typical experiment are presented

[b] Plasmid repair is defined as the number of $Kan^R$ colonies divided by the number of $Cm^R$ colonies. The average of six experiments is presented for cisPt-GG and three independent experiments for TT-CPD.

[c] TLS values are obtained by multiplying plasmid repair values by the fraction of TLS events, as deduced from the DNA sequence analysis.

**Appendix Table S5. Analysis of mutations formed during TLS across a cisPt-GG lesion in *TENT4A* or/and *TENT4B* knocked-down MCF-7 cells.**

| Event type | Sequence opposite 5'-A*GG*C-3' | Number of events | | | |
|---|---|---|---|---|---|
| | | siControl | *siTENT4A* | *siTENT4B* | *siTENT4A/B* |
| Accurate TLS | 3'-TCCG-5' | 56 | 51 | 52 | 52 |
| Mutagenic TLS | | | | | |
| Misinsertion opposite 3'of the lesion | 3'-TC**A**G-5' | 20 | 17 | 28 | 31 |
| | 3'-TC**T**G-5' | 2 | 4 | 2 | 2 |
| | 3'-TC**G**G-5' | 2 | 0 | 0 | 0 |
| Misinsertion opposite 5'of the lesion | 3'-T**A**CG-5' | 1 | 1 | 0 | 0 |
| | 3'-T**T**CG-5' | 0 | 0 | 1 | 0 |
| Semi-targeted misinsertion | 3'-TCC**T**-5' | 1 | 0 | 0 | 0 |
| | 3'-**A**CCG-5' | 0 | 0 | 1 | 1 |
| | 3'-TCC**G**-5' | 0 | 0 | 1 | 0 |
| | 3'-TCC**A**-5' | 0 | 0 | 1 | 0 |
| Double/triple mutation | 3'-**AA**CG-5' | 0 | 0 | 1 | 0 |
| | 3'-T**T**C**A**-5' | 0 | 1 | 0 | 0 |
| | 3'-T△△**A**-5' | 0 | 0 | 1 | 0 |
| | | | | | 0 |
| Tandem-double misinsetion | 3'-T**TA**G-5' | 1 | 0 | 0 | 0 |
| | 3'-T**AA**G-5' | 0 | 1 | 0 | 0 |
| | Total TLS events: | 83 | 75 | 88 | 86 |
| | Total mutagenic TLS events (%): | 27 (33%) | 24 (32%) | 36 (41%) | 34 (40%) |
| Non-TLS events | | 2 | 6 | 1 | 4 |
| Total events analyzed | | 85 | 81 | 89 | 90 |

Plasmids were extracted from Kan$^R$ colonies obtained in the experiments described in methods, and subjected to DNA sequence analysis. The cisPt-GG lesion is shown in italics (*GG*). Insertion of incorrect nucleotides is indicated by bold type letters. The two nucleotides incorporated opposite to the site of the cisPt-GG lesion are underlined. △ represents a single-nucleotide deletion. Semi-targeted misinsertion is a misinsertion event opposite a nucleotide flanking the lesion. Non-TLS events include big deletions and insertion of sequences from the control plasmid. The differences in the percent of mutagenic events among the various conditions did not reach statistical significance.

**Appendix Table S6. Analysis of mutations formed during TLS across a site-specific TT-CPD lesion in MCF-7 cells in which *TENT4A* expression was knocked-down.**

| Event type | Sequence opposite 5'-G*TT*G-3' | Number of events | |
|---|---|---|---|
| | | siControl | si*TENT4A* |
| Accurate TLS | 3'-CAAC-5' | 40 | 37 |
| Mutagenic TLS | | | |
| | 3'-C**T**AC-5' | 0 | 1 |
| | 3'-CA**G**C-5' | 0 | 1 |
| | 3'-**A**AAC-5' | 0 | 1 |
| | 3'-CAA**A**-5' | 0 | 1 |
| | 3'-CAA△-5' | 1 | 0 |
| | Total TLS events: | 41 | 41 |
| | Total mutagenic TLS events (%): | 1 (2.4%) | 4 (10%) |
| Non-TLS events | | 4 | 4 |
| Total events analyzed | | 45 | 45 |

Plasmids were extracted from $Kan^R$ colonies obtained in the experiments described in methods, and subjected to DNA sequence analysis. The TT-CPD lesion (cyclobutyl pyrimidine dimer) is shown in italics (*TT*). Insertion of incorrect nucleotides is indicated by bold type letters. The two nucleotides incorporated opposite to the site of the TT-CPD lesion are underlined. Non-TLS events include big deletions and insertion of sequences from the control plasmid. The differences in the percent of mutagenic events among the various conditions did not reach statistical significance (chi-squared test, *P*=0.166).

**Appendix Table S7. TLS across a site-specific cisPt-GG lesion in MCF-7 cells overexpressing *PAXIP1-AS2* long non-coding antisense RNA.**

| Sample | Lesion | Number of transformants[a] | | Plasmid repair[b], % | TLS[c], % | Relative TLS, % |
|---|---|---|---|---|---|---|
| | | $Kan^R$ | $Cm^R$ | | | |
| Empty vector (EV) | cisPt-GG | 122 | 219 | 54±2 | 52±2 | 100±4 |
| *PAXIP1-AS2* pCMV SPORT6[d] | cisPt-GG | 55 | 342 | 15±1 | 14±0.9 | 27±2 |

[a] The results of a typical experiment are presented

[b] Plasmid repair is defined as the number of $Kan^R$ colonies divided by the number of $Cm^R$ colonies. The average of two experiments is presented.

[c] TLS values are obtained by multiplying plasmid repair values by the fraction of TLS events, as deduced from the DNA sequence analysis.

[d] A plasmid overexpressing PAXIP-AS2 antisense RNA.

**Appendix Table S8. Analysis of mutations formed during TLS across a cisPt-GG lesion in MCF-7 cells overexpressing *PAXIP1-AS2* long non-coding RNA.**

| Event type | Sequence opposite 5'-A*GG*C-3' | Number of events | |
|---|---|---|---|
| | | Empty vector | *PAXIP1-AS2* Overexpressing cells |
| Accurate TLS | 3'-TCCG-5' | 38 | 37 |
| Mutagenic TLS | | | |
| | 3'-TC**A**G-5' | 4 | 1 |
| | 3'-TC**T**G-5' | 2 | 1 |
| | Total TLS events: | 44 | 39 |
| | Total mutagenic TLS events (%): | 6 (14%) | 2 (5%) |
| Non-TLS events | | 1 | 3 |
| Total events analyzed | | 45 | 42 |

Plasmids were extracted from $Kan^R$ colonies obtained in the experiments described in methods, and subjected to DNA sequence analysis. The cisPt-GG lesion is shown in italics (*GG*). Insertion of incorrect nucleotides is indicated by bold type letters. The two nucleotides incorporated opposite to the site of the cisPt-GG lesion are underlined. Non-TLS events include big deletions and insertion of sequences from the control plasmid. The differences in the percent of mutagenic events between the two conditions did not reach statistical significance.

**Appendix Table S9. TLS across a site-specific TT-CPD lesion in MCF-7 cells in which *PAXIP1* expression was knocked-down.**

| Sample | Lesion | Number of transformants[a] | | Plasmid repair[b], % | TLS[c], % | Relative TLS, % |
|---|---|---|---|---|---|---|
| | | $Kan^R$ | $Cm^R$ | | | |
| siControl | TT-CPD | 131 | 173 | 71±5 | 69±5 | 100±7 |
| si*PAXIP1* | TT-CPD | 45 | 149 | 35±5 | 34±4 | 50±6 |

[a] The results of a typical experiment are presented

[b] Plasmid repair is defined as the number of $Kan^R$ colonies divided by the number of $Cm^R$ colonies. The average of two experiments is presented.

[c] TLS values are obtained by multiplying plasmid repair values by the fraction of TLS events, as deduced from the DNA sequence analysis.

**Appendix Table S10. Analysis of mutations formed during TLS across a site-specific TT-CPD lesion in MCF-7 cells in which *PAXIP1* expression was knocked-down.**

| Event type | Sequence opposite 5'-G*TT*G3-3' | Number of events | |
|---|---|---|---|
| | | Empty vector | *PAXIP1-AS2* Overexpressing cells |
| Accurate TLS | 3'-C<u>AA</u>C-5' | 33 | 32 |
| Mutagenic TLS | | | |
| | 3'-C<u>**T**A</u>C-5' | 1 | 0 |
| | Total TLS events: | 34 | 32 |
| | Total mutagenic TLS events (%): | 1 (2.9%) | 0 (<3%) |
| Non-TLS events | | 1 | 1 |
| Total events analyzed | | 35 | 33 |

Plasmids were extracted from Kan$^R$ colonies obtained in the experiments described in methods, and subjected to DNA sequence analysis. The TT-CPD lesion (cyclobutyl pyrimidine dimer) is shown in italics (*TT*). Insertion of incorrect nucleotides is indicated by bold type letters. The two nucleotides incorporated opposite to the site of the TT-CPD lesion are underlined. Non-TLS events include big deletions and insertion of sequences from the control plasmid. The differences in the percent of mutagenic events between the two conditions did not reach statistical significance.

**Appendix Table S11. Number of samples with mutations in TLS-related *vs* random genes**[a]

| Cancer type[b] | Samples, n[c] | Samples with mutations, n | | *P*-total[f] |
|---|---|---|---|---|
| | | TLS-related genes[d] | Random genes, Ave[e] | |
| **LAML** | 144 | 21 | 5.029 | **<0.001** |
| **THYM** | 123 | 9 | 3.301 | **0.01** |
| **UCEC** | 530 | 196 | 153.918 | **0.029** |
| KIRP | 281 | 25 | 15.471 | 0.052 |
| SARC | 237 | 24 | 16.137 | 0.072 |
| HNSC | 508 | 84 | 57.257 | 0.077 |
| ESCA | 184 | 38 | 25.833 | 0.081 |
| THCA | 492 | 10 | 6.318 | 0.086 |
| PAAD | 180 | 11 | 7.018 | 0.089 |
| BRCA | 986 | 89 | 65.403 | 0.092 |
| UCS | 57 | 6 | 4.102 | 0.115 |
| CHOL | 51 | 5 | 3.35 | 0.127 |
| BLCA | 412 | 89 | 71 | 0.145 |
| DLBC | 37 | 6 | 4.238 | 0.146 |
| STAD | 437 | 106 | 87.165 | 0.152 |
| TGCT | 145 | 3 | 2.358 | 0.216 |
| COAD | 399 | 104 | 92.435 | 0.217 |
| LIHC | 364 | 38 | 33.199 | 0.261 |
| OV | 436 | 51 | 44.925 | 0.283 |
| KIRC | 336 | 19 | 16.59 | 0.289 |
| KICH | 66 | 2 | 1.966 | 0.307 |
| UVM | 80 | 1 | 1.122 | 0.315 |
| LUSC | 492 | 105 | 98.972 | 0.338 |
| CESC | 289 | 47 | 44.288 | 0.341 |
| MESO | 82 | 2 | 2.485 | 0.437 |
| PRAD | 496 | 13 | 13.92 | 0.467 |
| PCPG | 179 | 1 | 1.676 | 0.473 |
| READ | 137 | 18 | 19.872 | 0.527 |
| LGG | 509 | 11 | 14.001 | 0.635 |
| LUAD | 567 | 88 | 107.84 | 0.681 |
| GBM | 393 | 27 | 33.298 | 0.692 |
| SKCM | 468 | 121 | 144.477 | 0.717 |
| ACC | 92 | 3 | 5.935 | 0.843 |

[a] All mutations included.

[b] Cancer type according to TCGA cancer acronyms. https://gdc.cancer.gov/resources-tcga-users/tcga-code-tables/tcga-study-abbreviations

[c] Number of samples for which sequence information is available in the TCGA database.

[d] Number of samples with a mutation in at least one of 14 TLS-related genes listed in Table 1b.

[e] Number of samples with a mutation in at least one of 14 random genes; Average of 1000 random sets.

[f] Fraction of the 1000 random runs which yielded a number of samples ≥ fraction of TLS genes.

See also Appendix Figure S4.

**Appendix Table S12. siRNA used for the knockdown studies**

| Gene Name | Catalog Number | Full Name | Sequence (5'-3') |
|---|---|---|---|
| *TENT4A* | M-009807-01 | SMARTpool: siGENOME TENT4A siRNA | GGAGUGACGUUGAUUCAGA |
| | | | CAGACAAGCUGGUGUAGAA |
| | | | GCCCUCAUCUGUAUCAUAA |
| | | | GCCAUUAGCUUUCUACAGU |
| *TENT4B* | M-010011-01 | SMARTpool: siGENOME TENT4B siRNA | GGACGACACUUCAAUUAUU |
| | | | GAUAAAGGAUGGUGGUUCA |
| | | | GAAUAGACCUGAGCCUUCA |
| | | | UAUCGAAGAUCCUUUACAA |
| *CYLD* | M-004609-01 | SMARTpool: siGENOME CYLD siRNA | CGAAGAGGCUGAAUCAUAA |
| | | | GAACAGAUUCCACUCUUUA |
| | | | GAACUCACAUGGUCUAGAA |
| | | | GGACAUGGAUAACCCUAUU |
| *DTL* | M-020543-00 | SMARTpool: siGENOME DTL siRNA | ACUCCUACGUUCUCUAUUA |
| | | | GUAUGGGAUUUACGUAAGA |
| | | | AGAAGGCUUUGUUCGAUUG |
| | | | GCUAAUUGCACAGACGAUA |
| *RAD18* | L-004591-00 | SMARTpool: ON-TARGET*plus* RAD18 siRNA | CCAAGAAACAAGCGUAAUA |
| | | | GGGAGCAGGUUAAUGGAUA |
| | | | GCUCUCUGAUCGUGAUUUA |
| | | | GAAAUGAGUGGUUCUACAU |
| *PAXIP1* | | *PAXIP1 (Gohler et al, 2008)* | UGCACUAGCCUCACACAUA |
| Non-targeting control#5 | D-001210-05 | siGENOME Non-Targeting Control siRNA #5 | UAGCGACUAAACACAUCAA |
| Non-targeting control pool | D-001810-10 | ON-TARGET*plus* Non-targeting Pool | UGGUUUACAUGUCGACUAA |
| | | | UGGUUUACAUGUUGUGUGA |
| | | | UGGUUUACAUGUUUUCUGA |
| | | | UGGUUUACAUGUUUUCCUA |

**Appendix Table S13. List of qPCR primers used in this study**

| Gene | Forward primer (5'→3') | Reverse primer (5'→3') |
|---|---|---|
| *POLH* | ATGATCCAGTTAAACCCAGG | CATTTCGGTCTTTAGTCAGTC |
| *POLK* | AGGTTGACAGATTTGCAATG | AGAAAGCATCCATGTCAATG |
| *POLI* | GGAAATTATGATGTGATGACCC | TCAATAAGCCCTTTCTTAGC |
| *REV1* | CAGACATCAGAGCTGTATAATG | CCTTTGAGATCTGGTCTATTTC |
| *REV3L* | AGGTCACACTAGACACAAAG | GTCAAAGGCTGAAAAGTCTG |
| *RAD18* | AAAGTGGATTGTCCTGTTTG | CTTCTCTTCGCGTGATAAAC |
| *PCNA* | CTGTGTAGTAAAGATGCCTTC | TCTCTATGGTAACAGCTTCC |
| *CYLD* | CTACAGGATCTACCTCAGAC | GTCAACAGAAGACCCATTTAAG |
| *PAXIP1* | AGTTCCTGACGGCGATTTCTGT | AAGTACTTCTGCCTCAGCATCTCG |
| *PAXIP1-AS2* | GCTTCCAGGGGAGAT | ATCAGACTGCCTGAAGA |
| *RNA18S* | ACAGGATTGACAGATTGA | TATCGGAATTAACCAGACA |
| *GAPDH* | ACAGTTGCCATGTAGACC | TTTTTGGTTGAGCACAGG |
| *TENT4A* | TGAAGTGAAAGTTGACAT | GTAAGGCAGCAATGAATA |
| *TENT4B* | CCAACAATGAAACAGAAAGC | AAGGCTCAGGTCTATTCTTC |
| *POLD3* | GAAGCGAGTAGCATTATCTG | GACTTCATCTTCACTGCTATC |

**Appendix Table S14. List of  primers used in the ePAT assay**

| | Primer (5'-3') |
|---|---|
| *RAD18* | CCTGTCTTCCCCAAAATTCA |
| *CYLD* | CTGCATTTCCCATTTTTGGT |
| *POLH* | TCACGTGAAAGGCCTCAGTA |
| TVN-PAT | GCGAGCTCCGCGGCCGCGTTTTTTTTTTTTVN |
| ePAT-Anchor | GCGAGCTCCGCGGCCGCGTTTTTTTTTTTT |

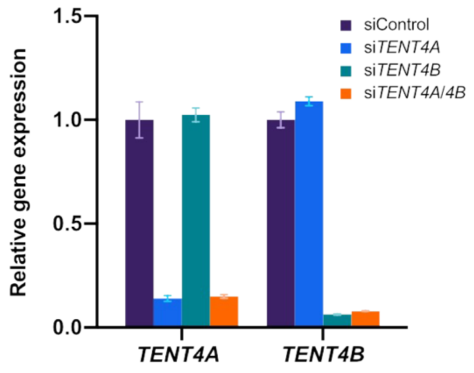


**Appendix Figure S1.** Knockdown of *TENT4A* and/or *TENT4B* in MCF-7 cells. Expression of *TENT4A* and *TENT4B* alone or in combinations were knocked-down for 48 h, after which the level of *TENT4A and TENT4B* transcripts were measured by qPCR, normalized to *GAPDH*, and compared to cells treated with siControl. The results are presented as the mean ± SEM of three independent experiments.

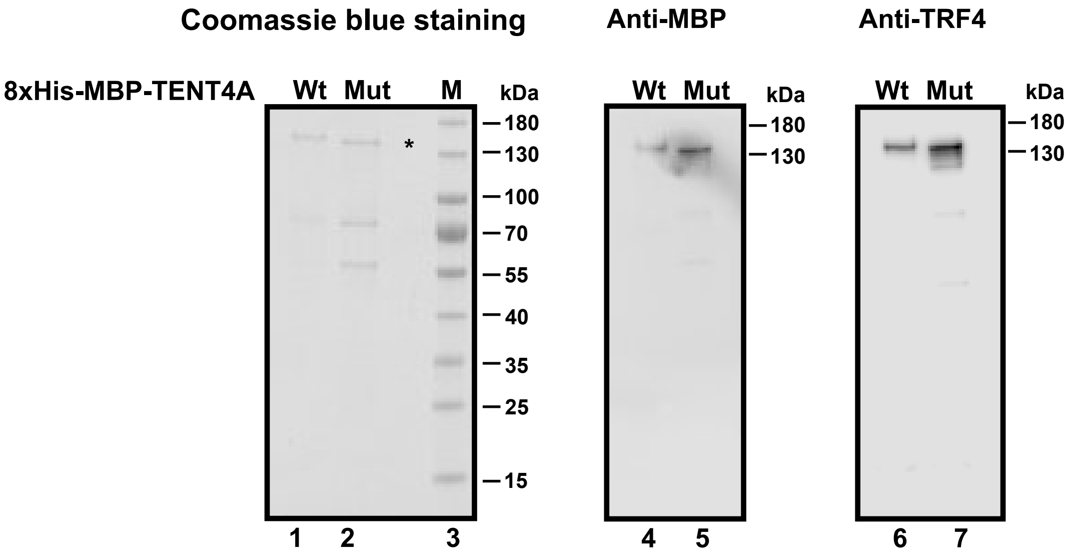


**Appendix Figure S2**. Partially purified 8xHis-MBP-tagged TENT4A (Wt) and mutant 8xHis-MBP-tagged TENT4A DD277,279AA (Mut) in which Aspartate at positions 277 and 279 have been changed to Alanine, were analyzed on a 4-20 % gradient sodium dodecyl sulphate-polyacrylamide gel and stained with Coomassie blue. The position of the partially purified proteins is indicated by an asterisk. Lane 1, 0.5 µg of Wt protein; lane 2, 0.5 µg of Mut protein; lane 3, molecular mass standards.

Wt and mutant proteins are analyzed by western blotting using the indicated antibodies.

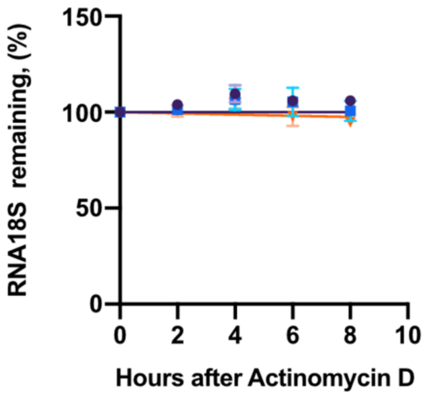


**Appendix Figure S3**. Effect of TENT4A on the stability of RNA18S rRNA. Knocking down the expression of *TENT4A*, *TENT4B* or both, had no detectable effect on the stability of RNA18S rRNA. Data are presented as mean ± SEM of three independent experiments.

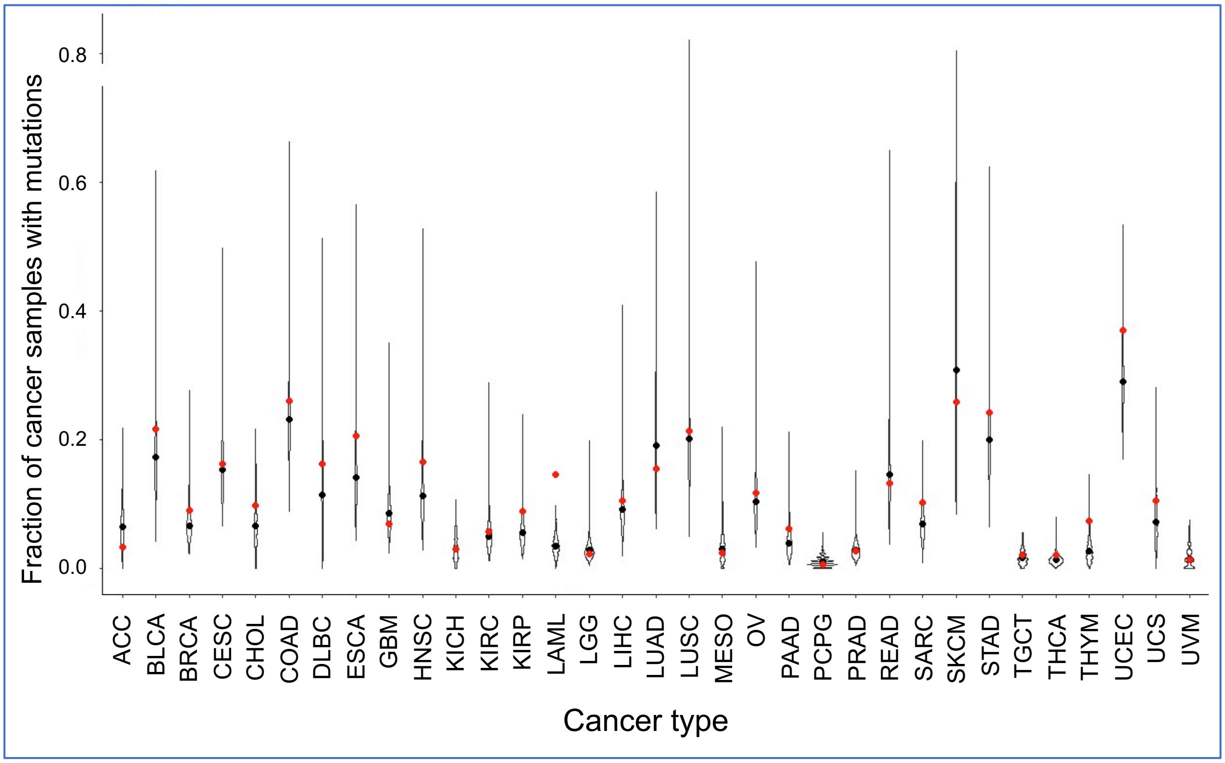


**Appendix Figure S4.** Distribution of the fraction of samples with mutations in sets of random 14 genes. The distribution of 1000 such independent sets is shown for each cancer type. The red dot marks the fraction of samples with mutations in at least one gene of the 14 TLS-related genes *CYLD*, *NPM1*, *TENT4B*, *PAXIP1*, *PCNA*, *POLH*, *POLI*, *POLK*, *PRIMPOL*, *RAD18*, *REV1*, *REV3L,* and *USP1.* The probability that the fraction of samples with mutations in TLS-related genes was obtained by chance is <0.05 only for 3 cancer types: LAML, THYM, and UCEC (see Appendix Table S11).
